# *In vitro* characterization of the human segmentation clock

**DOI:** 10.1101/461822

**Authors:** Margarete Diaz-Cuadros, Daniel E Wagner, Christoph Budjan, Alexis Hubaud, Jonathan Touboul, Arthur Michaut, Ziad Al Tanoury, Kumiko Yoshioka-Kobayashi, Yusuke Niino, Ryoichiro Kageyama, Atsushi Miyawaki, Olivier Pourquié

**Affiliations:** Department of Genetics, Harvard Medical School and Department of Pathology, Brigham and Women’s Hospital, Boston, MA, USA; Department of Systems Biology, Harvard Medical School, Boston, MA, USA; Department of Mathematics, Brandeis University, MA, USA; Institute for Frontier Life and Medical Sciences, Kyoto University, Kyoto, Japan; Laboratory for Cell Function and Dynamics, Brain Science Institute Riken, Saitama, Japan; Harvard Stem Cell Institute, Harvard University, Cambridge, MA USA

## Abstract

The vertebral column is characterized by the periodic arrangement of vertebrae along the anterior-posterior (AP) axis. This segmental or metameric organization is established early in embryogenesis when pairs of embryonic segments called somites are rhythmically produced by the presomitic mesoderm (PSM). The tempo of somite formation is controlled by a molecular oscillator known as the segmentation clock ^1,2^. While this oscillator has been well characterized in model organisms ^1,2^, whether a similar oscillator exists in humans remains unknown. We have previously shown that human embryonic stem (ES) cells or induced pluripotent stem (iPS) cells can differentiate *in vitro* into PSM upon activation of the Wnt signaling pathway combined with BMP inhibition^3^. Here, we show that these human PSM cells exhibit Notch and YAP-dependent oscillations^4^ of the cyclic gene *HES7* with a 5-hour period. Single cell RNA-sequencing comparison of the differentiating iPS cells with mouse PSM reveals that human PSM cells follow a similar differentiation path and exhibit a remarkably coordinated differentiation sequence. We also demonstrate that FGF signaling controls the phase and period of the oscillator. This contrasts with classical segmentation models such as the “Clock and Wavefront” ^1,2,5^, where FGF merely implements a signaling threshold specifying where oscillations stop. Overall, our work identifying the human segmentation clock represents an important breakthrough for human developmental biology.

---

Twenty years ago, the segmental organization of the vertebral column in the chicken embryo was shown to be controlled by a molecular oscillator called “Segmentation clock” ^6^. This oscillator was subsequently identified in mouse, frog and zebrafish suggesting that it represents a conserved feature of vertebrate segmentation^1,2^. The clock controls the periodic activation of the Notch, Wnt and FGF signaling pathways which drive the rhythmic production of embryonic somites from the PSM^1,2^. Analyses of consanguineous families with probands exhibiting severe spine segmentation defects (congenital scoliosis) have implicated several human orthologs of cyclic genes associated with the mouse segmentation clock, such as *HES7*^7^ or *LUNATIC FRINGE*^8^, suggesting that this oscillator might be conserved in humans. A total of 38-39 somites form between 20 and 30 days post-fertilization in humans^9^, resulting in the production of the 33 vertebrae of adults. However, the somitogenesis stages of human embryos are very difficult to access and the impossibility to culture post-implantation embryos means that virtually nothing is known about human segmentation beyond histological descriptions^9^.

We have previously shown that the early stages of human paraxial mesoderm development can be recapitulated *in vitro* by treating iPS cells with Wnt activators such as the GSK3β inhibitor CHIRON99021 (Chir) in combination with the BMP inhibitor LDN193189 (LDN) (CL medium)^3,10^ (Fig. 1a). Following one day of CL treatment, iPS cells first transition from pluripotency to a state resembling neuro-mesodermal progenitors (NMPs)^11,12^/anterior primitive streak as suggested by the expression of both *T/BRACHYURY*, *SOX2* and *POU5F1 (OCT4)*(Fig 1a-b, Extended Data Fig. 1a). By day 2 of differentiation, these cells activate the paraxial mesoderm markers *TBX6* and *MSGN1* (Fig. 1a). At the same time, an epithelium-to-mesenchyme transition (EMT) marked by a switch from *CDH1* to *CDH2* takes place and is accompanied by active migration of cells away from epithelial colonies as *MSGN1* begins to be expressed (Extended Data Fig. 1a; Movie S1). This process strikingly parallels cell ingression in the primitive streak of amniote embryos. Paraxial mesoderm induction under CL conditions is highly efficient, as differentiation of an iPS line harboring a *MSGN1-Venus* knock-in reporter construct^13^ in CL medium shows that the reporter is activated in more than 95% of the cells from day 2 to 4 (Extended Data Fig. 1b).

**Figure 1.**
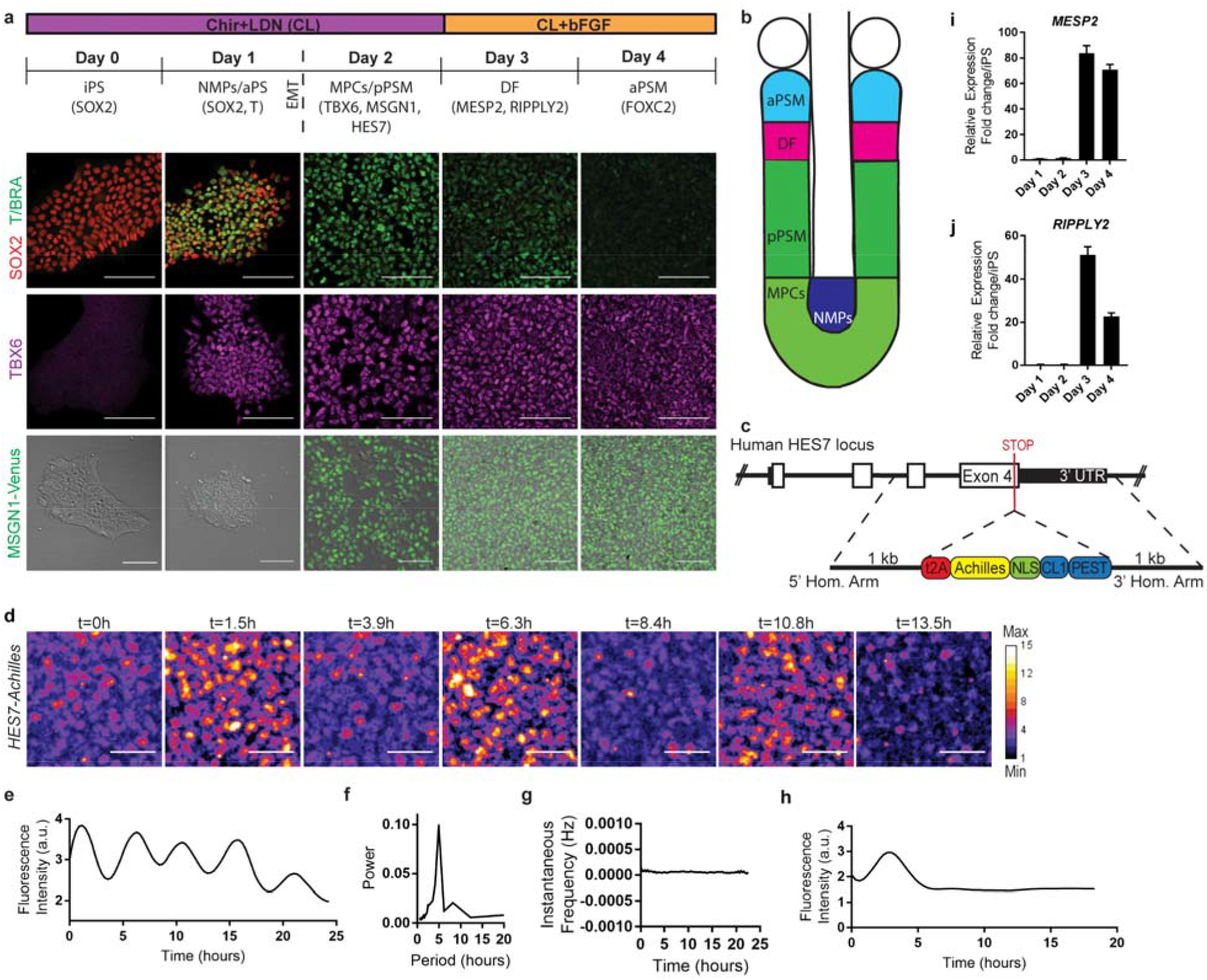
A *HES7-Achilles* fluorescent reporter oscillates with a 5-hour period in human PSM-like cells differentiated *in vitro*. **a,** Differentiation timeline outlining major changes in cell fate and medium conditions. iPS, induced pluripotent stem cells; aPS, anterior primitive streak; NMPs, neuromesodermal progenitors; MPCs, mesodermal precursor cells; pPSM, posterior presomitic mesoderm; aPSM, anterior presomitic mesoderm; DF, determination front. Top: Immunofluorescence staining for T/BRACHYURY and SOX2 on days 0-4 of differentiation. Middle: Immunofluorescence staining for TBX6 on days 0-4 of differentiation. Bottom: Brightfield and fluorescence merged images of the MSGN1-Venus reporter line on days 0-4 of differentiation. Scale bar = 100μm **b,** Scheme illustrating the maturation stages of paraxial mesoderm. aPSM, anterior PSM; DF, determination front; MPCs, mesodermal precursor cells; pPSM, posterior PSM; NMPs, neuromesodermal progenitors. **c,** Diagram outlining the targeting strategy used to generate a *HES7-Achilles* knock-in reporter line in human iPS cells. **d,** Snapshots of *HES7-Achilles* fluorescence showing peaks and troughs over the course of 13.5 hours. Scale bar = 100μm **e,** Quantification of *HES7-Achilles* fluorescence intensity from day 2 to day 3 of differentiation. **f,** Fourier transform of *HES7-Achilles* oscillations indicating the predominant period. **g,** Instantaneous frequency in Hertz (calculated by Hilbert transformation) of *HES7-Achilles* oscillations over time **h,** Quantification of *HES7-Achilles* fluorescence over the course of 20 hours in cells differentiated without the BMP inhibitor LDN93189 (CHIR99021 only medium). **i,** qRT-PCR for the determination front marker *MESP2* on days 1-4 of differentiation. Relative expression expressed as fold change relative to iPS (day 0). Mean ±SD **j,** qRT-PCR for the determination front marker *RIPPLY2* on days 1-4 of differentiation. Relative expression expressed as fold change relative to iPS (day 0). Mean ±SD

Expression of segmentation clock genes such as *HES7* and *LFNG* has been described in differentiating human iPS cultures during this time window, but their expression dynamics has not been investigated ^3,13,14^(Extended Data Fig. 1c). In mouse, oscillations of a *Hes7-Luciferase* reporter in ES cells differentiated *in vitro* to a PSM fate have been recently reported ^15^. To visualize oscillations of the human segmentation clock, we used CRISPR-Cas9 technology to knock-in a destabilized version of the rapidly folding YFP variant Achilles into the 3’ end of *HES7* in a human iPSC line ^16^(Fig. 1c). A 2A peptide linker was used to ensure cleavage of the fluorescent protein from the endogenous HES7 protein during translation. When the *HES7-Achilles* iPS line was differentiated in CL medium and imaged using time-lapse confocal microscopy, the majority of cells underwent a series of up to five oscillations between day 2 and day 3 (Fig. 1d-e, Movie S2). In these conditions, we did not observe traveling waves as described in mouse explants or in differentiated ES cells^4,15,17,20^. The period of oscillations was close to 5 hours and their frequency was constant over time (Fig. 1f-g). Interestingly, this period is similar to that reported for *HES1* oscillations in human cells derived from umbilical cord blood exposed to a serum shock^18^. No oscillations could be observed when LDN was omitted from the culture medium (Fig. 1h), supporting the notion that BMP inhibition is required to maintain PSM identity ^13^. *In vivo*, PSM cells stop oscillating and respond to the clock signal when they reach the determination front, a level defined by thresholds of FGF and Wnt signaling that form posterior to anterior gradients along the PSM^1^. In response to the clock signal, competent cells activate the expression of genes such as *Mesp2,* which defines the anterior and posterior boundaries of the future segment^19^. We observed that the arrest of *HES7-Achilles* oscillations coincided with the onset of expression of *MESP2* and *RIPPLY2* at day 3 as observed in mouse *in vivo* (Fig. 1i-j). Together, these data demonstrate that human iPS cells differentiated to a posterior PSM fate recapitulate oscillations of the segmentation clock followed by segmental patterning.

To further characterize the identity of the human PSM cells generated *in vitro*, we compared their transcriptomes to those of the *in vivo* mouse PSM cells. Using the inDrops single cell RNA-sequencing (scRNA-seq) platform^20^, we analyzed 5,646 cells dissociated from the entire posterior region including the two PSMs and the last three somites, the tail bud and the neural tube of two E9.5 mouse embryos. Clustering analysis revealed 19 distinct cell states corresponding to expected derivatives of posterior mesoderm including the PSM, and the endoderm and ectoderm-derived tissues (Fig. 2a, Extended Data Fig. 2a-d and Table S1). We next performed a detailed analysis of cells from the PSM and the neural tube clusters (Fig. 2b), which share a common developmental origin ^11,21^. When isolated and represented as a k-nearest neighbor (kNN) graph, these cells organized into a continuum of states recapitulating spatio-temporal features of the developing vertebrate tailbud. Identification of differentially expressed genes along a pseudo-temporal trajectory (Fig. 2c and Table S2) and between subclusters of cells (Fig 2d, f) revealed distinct phases of paraxial mesoderm differentiation, arranged sequentially on the knn graph. One cluster, characterized by expression of *Sox2* and T/*Brachyury* (the NMP expression signature) was positioned at an intermediate location in the overall continuum and was flanked by clusters for posterior neural tube and paraxial mesoderm, respectively. This ordering of states is consistent with the known bipotentiality of NMPs for these lineages ^11^. Several additional clusters describe successive stages of PM differentiation from NMPs (Extended Data Fig. 3a-c and Table S3). Two clusters define progressively the most immature PSM cells, which retain T expression and are called mesodermal precursor cells (MPCs)^22^, and more mature posterior PSM (pPSM) (Fig. 1b). These two clusters define the oscillatory domain *in vivo*. The determination front, highlighted by *Mesp2* expression, marks the boundary of the next cluster, the anterior PSM (aPSM), which is characterized by the onset of *Pax3* expression. Two additional clusters correspond to the ventral (sclerotome) and dorsal (dermomyotome) regions of the newly formed somites. This inferred organization of cell states was reproducible by an independent clustering approach (Extended Data Fig. 3a-b).

**Figure 2.**
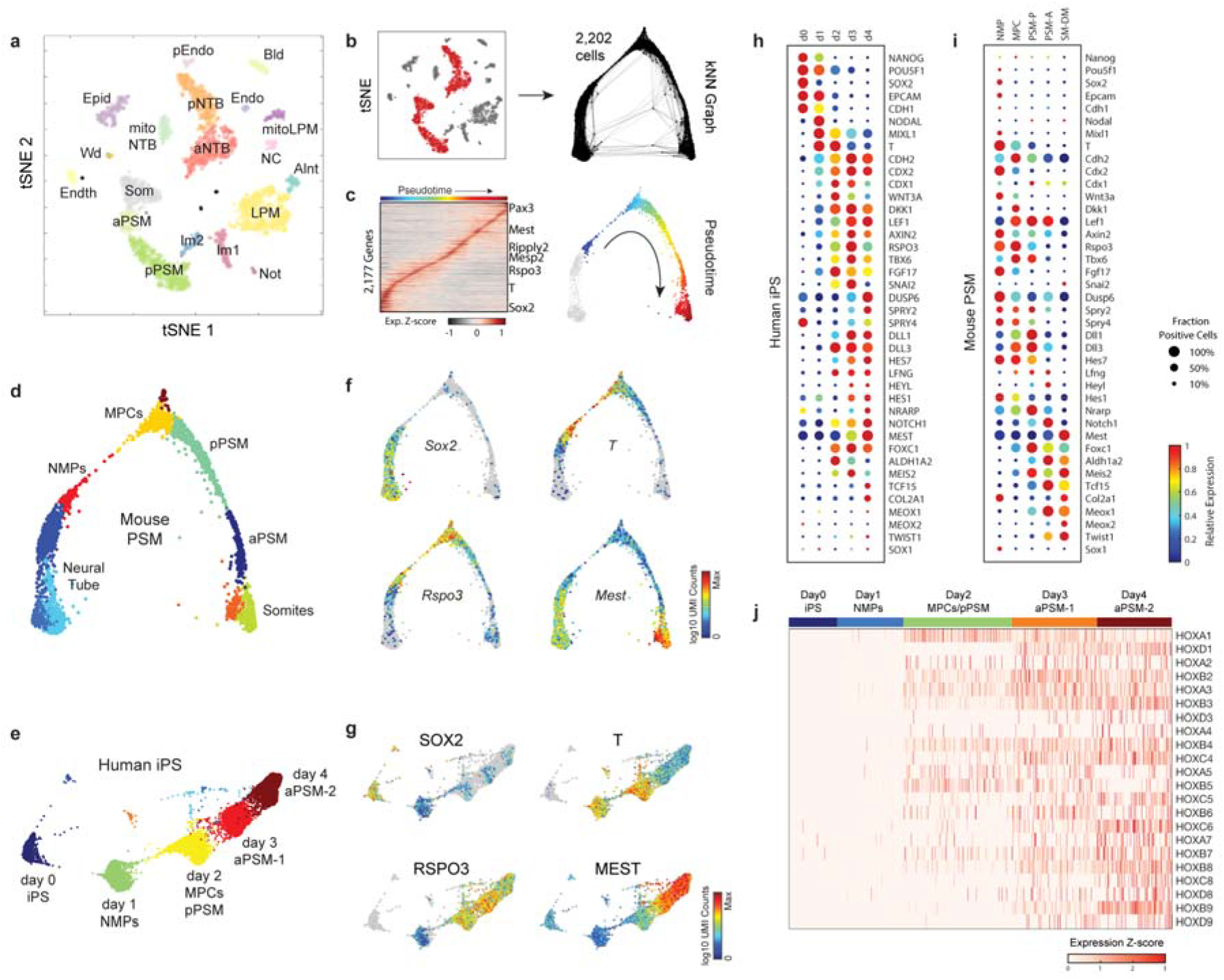
Single cell RNA-sequencing analysis of differentiating mouse and human PSM. **a,** tSNE embedding of E9.5 cells, colored by density cluster ID (see also Extended data Fig.2c-d and Table S1). **b,** Clusters corresponding to neural tube, presomitic, and somites were isolated (left, red dots), and used to construct a k-nearest-neighbor (kNN) graph (right, 2,202 cells, k = 20, 12 PC dimensions). **c,** Pseudotemporal ordering of non-neural cells and identification of dynamically expressed genes. Heatmap illustrates genes with significant dynamic expression, ordered by peak expression. Select genes marking progressive stages of paraxial mesoderm differentiation are labeled. **d,** E9.5 kNN graph with cell nodes colored by Louvain cluster ID (see Fig S2a-c and Table S3). **e,** kNN graph of human iPS cultured cells (14,750 cells, k = 30, 17 PC dimensions), colored by Louvain cluster ID (see also Extended Data Figure 3d-f and Table S4). **f-g,** Mouse E9.5 and human iPS kNN graph nodes colored by expression levels of orthologous genes. **h-i,** Dot plots illustrating single-cell expression characteristics for select genes for each human iPS collection timepoint (h) or mouse E9.5 cluster (i). Colors indicate expression level, dot sizes indicate fraction of positive cells in each human timepoint or E9.5 cluster, respectively. **j,** Heatmap of expression levels for human HOX genes (>5 transcripts per million, TPM). Rows (individual cells) are ordered by timepoint, columns (HOX genes) are ordered by position.

We next compared the progression of these *in vivo* mouse cell states to those of 14,750 human iPS-derived cells analyzed by scRNA-seq over the first 4 days of *in vitro* PSM differentiation. Similar to above, a knn-graph-based clustering analysis revealed an ordered progression of cell states (Fig. 2e). The vast majority of single-cell transcriptomes from these iPS-derived cultures (>93%) comprised 5 major clusters expressing markers of increasingly differentiated cells (Fig. 2g, Extended Data Fig. 3d-f and 4, Table S4). Strikingly, this unsupervised analysis organized collection timepoints sequentially along the knn graph, each of which was dominated by a single cluster (Fig. 2e, Extended Data Fig. 3d-f). Differential gene expression analysis revealed shared molecular characteristics between the human clusters and the *in vivo* mouse paraxial mesoderm lineage (Fig. 2h-i, Extended Data Fig. 3c, f). For example, cells collected after day 1 of differentiation exhibited characteristics of NMPs/anterior primitive streak with expression of genes such as *NODAL*, *T, MIXL1* and *SOX2* (Fig. 2h-i, Extended Data Fig 4a). Day 2 human cells resembled the mouse MPC and posterior PSM clusters with expression of genes such as *T, MSGN1, TBX6, DLL3, WNT3A* and *FGF17*, as well as the Notch-associated cyclic genes *LFNG* and *HES7* (Fig. 2h-i, Extended Data Fig 4a). Day 3 and Day 4 cells show expression of markers of more anterior PSM such as *RSPO3*, *MEST* and *FOXC1* (Fig. 2h-I, Extended Data Fig 4a). Remarkably, we could also detect the collinear activation of HOX genes (Fig. 2j). Paralogs from all four HOX clusters were progressively expressed in a 3’ to 5’ order starting with *HOXA1* and *HOXA3* as early as day 1 while expression of *HOXB9* and *C8* peaked at day 4 (Extended Data Fig 4b). Genes of the paralog groups 10 to 13 were not detected arguing that the paraxial mesoderm cells produced exhibit a thoracic identity. Together, these analyses demonstrate that differentiating human iPS cells to a PSM fate *in vitro* in CL medium recapitulates a developmental sequence similar to that of the mouse embryo leading to the production of trunk paraxial mesoderm cells. Moreover, these data show that differentiation proceeds in a remarkably synchronized fashion resulting in strikingly homogenous populations in these cultures.

We recently demonstrated that the total number of oscillations can be increased in mouse tailbud explants by culturing in CL medium supplemented with Fgf4, the RA inhibitor BMS493 and the Rho kinase inhibitor (ROCKi) Y-27362 (CLFBR medium) ^4^ (Fig. 3a). When human *HES7-Achilles* reporter cells were cultured in such conditions from day 2 onwards, no dampening of the oscillations was observed and up to ten oscillations of the reporter were detected during day 2 and 3 (Fig. 3b, Movie S3). These oscillations retained the same 5 hour-period and their frequency was stable (Extended Data Fig. 5a-c). The treatment did not affect the percentage of MSGN1-Venus positive cells between day 2 and 4 (Extended Fig. 5d). Extension of the oscillatory window was accompanied by a delay in the onset of expression of determination front genes (*MESP2* and *RIPPLY2*) as well as anterior PSM markers (*FOXC2*) from day 3 to day 4 (Extended Data Fig. 5e). These new conditions provided us with an optimal system to investigate the regulation and the dynamical properties of the human segmentation clock oscillator.

**Figure 3.**
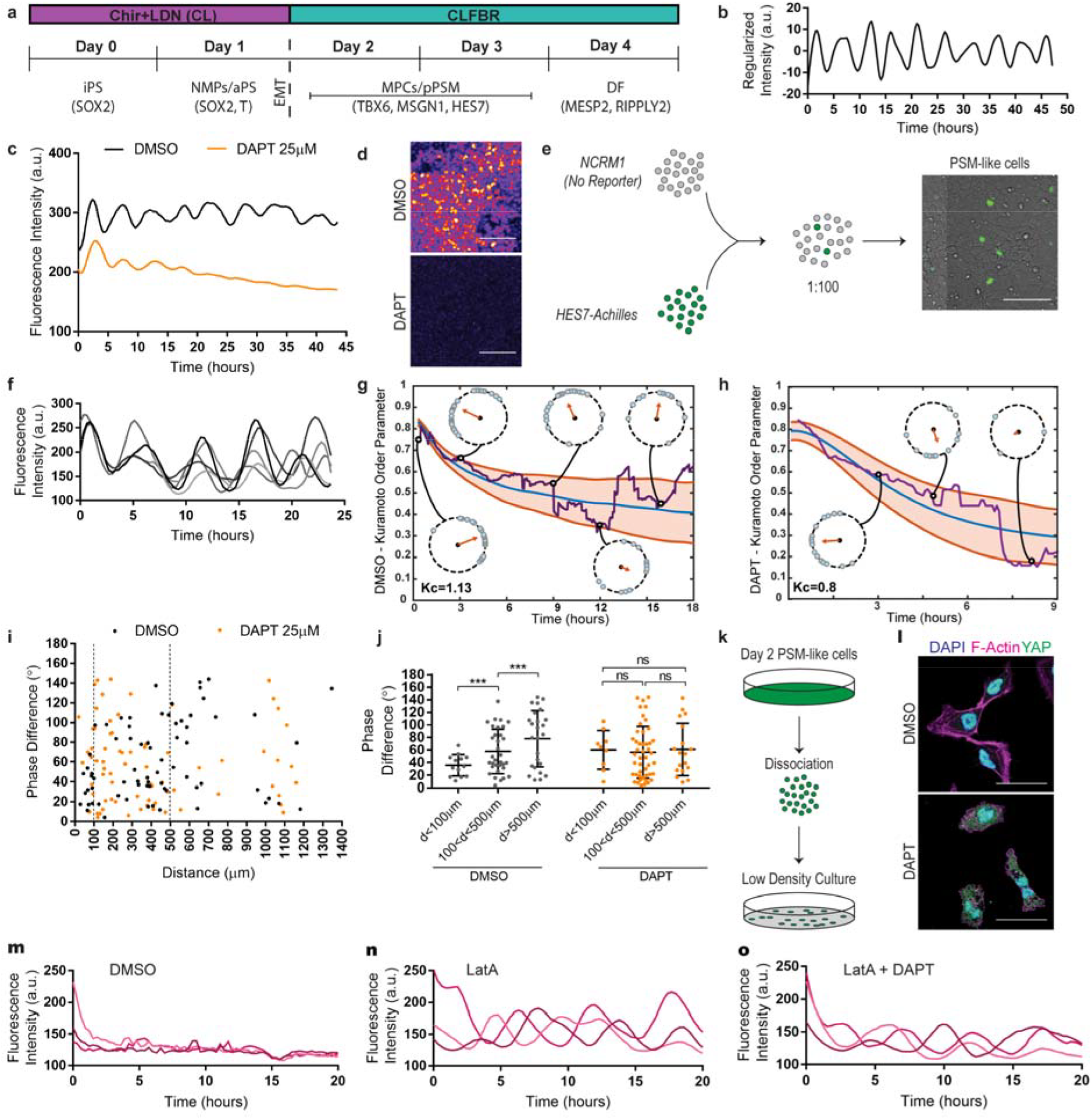
Notch and YAP signaling control oscillations of the *HES7-Achilles* reporter. **a,** Timeline outlining modified differentiation protocol for extension of the posterior PSM fate and increase in the HES7-Achilles oscillatory window. Major changes in cell fate and medium conditions used are indicated. iPS, induced pluripotent stem cells; aPS, anterior primitive streak; NMPs, neuromesodermal progenitors; MPCs, mesodermal precursor cells; pPSM, posterior presomitic mesoderm; DF, determination front. **b,** *HES7-Achilles* fluorescence intensity (regularized to oscillate about the origin by removing the moving average over 10 timepoints) over the course of 48 hours (Days 2-4 of differentiation) in CLFBR conditions. **c,** *HES7-Achilles* fluorescence intensity over a period of 45 hours in cells treated with DMSO (vehicle control) or the γ-secretase inhibitor DAPT (25μM) in CLFBR medium. **d,** Images of *HES7-Achilles* fluorescence following treatment with DMSO or DAPT (25μM). Scale bar = 100μm**e,** Outline of the experimental strategy for the tracking of *HES7-Achilles* oscillations in individual cells. Scale bar = 100μm **f,** *HES7-Achilles* fluorescence intensity in individual cells tracked over a period of 24 hours. Each line represents a different cell. All cells shown were initially located <100 μm away from each other. **g-h,** Evolution of the Kuramoto order parameter over time (purple line) for either DMSO (g) or DAPT-treated (25μM; h) cells, fitted with a Kuramoto model (blue line ±SD orange lines). Circles show the phase distribution of oscillators and their average vector (arrow) at different time points. Coupling constant (Kc) values are shown. **i,** Graph showing the phase difference of *HES7-Achilles* oscillations (degrees) between individual cells as a function of the initial distance between the cells (μm) under control (DMSO-black) and DAPT (25μM-orange) conditions. **j,** Mean ±SD phase difference (degrees) for *HES7-Achilles* oscillations between individual cells under control (DMSO-gray) and DAPT (25μM-orange) conditions. Same data as Figure 3i but binned by distance. *p≤0.001 (t-test) **k,** Diagram outlining the dissociation of PSM-like cells (Day 2) and reseeding at low density in fibronectin-coated dishes for the analysis of oscillations in isolated cells (i.e. no cell-cell contacts). **l,** Immunofluorescence staining for YAP, F-actin (phalloidin) and nuclear stain in isolated PSM-like cells treated with either DMSO or Latrunculin A (350nM). Scale bar = 50 μm. **m-o**, *HES7-Achilles* fluorescence intensity profiles for three different isolated cells individually tracked over the course of 20 hours. Cells were dissociated on Day 2 of differentiation and reseeded at low density on fibronectin-coated plates in CLFBR medium containing either DMSO (m), 350nM Latrunculin A (n), or 350nM Latrunculin A in combination with 25μM DAPT (o).

We first set out to characterize the role of Notch signaling in the regulation of *HES7-Achilles* oscillations. In mouse and zebrafish, Notch has been implicated in both the onset of oscillations and their local synchronization ^4, 23-25^. Treatment of *HES7-Achilles* cells with the Notch inhibitor DAPT in CLFBR medium on day 2 led to a rapid dampening of oscillations resulting in a decrease of target genes and reporter expression (Fig. 3c-d; Extended Data Fig. 6a, Movie S4). This indicates that *HES7* oscillations require active Notch signaling as recently reported for both *Lfng* and *Hes7* oscillations in mouse tailbud explants and mESC-derived PSM cells^7,12^. To test whether Notch also mediates the synchronization of individual oscillators, we attempted to track oscillations in single cells. Oscillating cells are however extremely motile as expected for posterior PSM cells ^26^, making the tracking of single cells difficult in these cultures. In fact, the average diffusivity of individual cells tracked *in vitro* (1.6±0.7 μm^2^/min; Fig. Extended Data Fig. 6b) is comparable to the diffusivity of PSM cells in the chicken embryo, which ranges from 0.5-8μm^2^/min *in vivo* ^19^. To quantitatively analyze the synchronization between individual cells in oscillating cultures, we diluted the cells carrying the *HES7-Achilles* reporter with their parental line (no reporter) in a ratio of 1:100 (Fig. 3e). As a result, Achilles-positive cells were sparse within the culture allowing individual cells to be tracked (Movie S5). DAPT-treated cells could not be tracked for longer than 10 hours on account of severe dampening. At the population level, cells are generally well synchronized in the control condition (Fig. 3f; Extended Data Fig. 6c), with an average phase shift of 36° (sem± 5.63°; n=29). We estimated an instantaneous phase shift through local cross-correlation, and compared the phase distribution with the classical Kuramoto model ^27^. The data was found consistent with the synchronized regime (coupling 13% larger than critical threshold, see Fig. 3g) in control conditions, but consistent with a Kuramoto model within the disordered region (with coupling parameter 20% smaller than critical coupling, see Fig. 3h) in the case of DAPT treatment. Furthermore, synchronization between control cells was stronger locally, and progressively became looser (typical distance of 200μm, see Extended Data Fig. 6e and Fig. 3i-j, linear fit R^2^=0.21, F-statistics vs constant model p<10^-3^), with close neighbors showing a higher degree of synchronization than remote cells. Pairwise phase of nearby cells (<100 μm apart, n=13) show an average phase shift twice smaller than cells at intermediate distances (>100μm and <500μm apart, n=33, 2-sample t-test p=0.007) and three times smaller than remote cells (>500μm, n=19, 2-sample t-test p<10^-3^ [nearby cells] and p=0.011 [intermediate cells], Fig. 3i-j, Ext. Data Fig. 6d). DAPT treated cells showed no such local synchrony, as no difference was found in the average phase shift of nearby cells compared to remote cells (Fig. 3i-j). These results were confirmed using various definitions of instantaneous phases (Extended Data Fig. 6f-j). Thus, loss of Notch signaling seems to abolish local coupling, leading to desynchronization in the cycles prior to oscillatory arrest.

We further assessed whether oscillations in human cells are regulated by YAP-mediated mechanical cues as recently demonstrated for mouse PSM cells cultured *ex vivo^4^*. Indeed, when human cells were dissociated on day 2 of differentiation and reseeded as isolated cells on fibronectin in CLFBR medium, no oscillations were detected (Fig. 3m, Movie S6). However, when treated with the F-actin inhibitor Latrunculin A, known to inhibit YAP ^28^, oscillations of normal period were restored (Fig. 3n, Movie S6). Even when Notch signaling was blocked with the γ-secretase inhibitor DAPT, isolated cells treated with Latrunculin A continued to oscillate (Fig. 3o, Movie S6). Our previous work suggested that the mouse segmentation clock exhibits dynamic properties of an excitable system where Notch is involved in triggering excitation while YAP controls the excitability threshold^4^. We now demonstrate that the human segmentation clock is regulated by Notch and YAP signaling in a similar fashion to the mouse clock arguing that excitable properties are conserved features of the mammalian oscillator.

*In vivo*, posterior-to-anterior gradients of FGF and Wnt signaling control PSM maturation^1^. We therefore examined the regulation of FGF and Wnt signaling in differentiating iPS cultures. On day 1 and 2, differentiating iPS cells exhibited strong immunofluorescence staining for doubly phosphorylated ERK (dpERK), which was down regulated on day 3 (Fig. 4a). The dpERK signal observed on day 2 was down-regulated by treatment with the FGF receptor inhibitor PD173074 (PD17), suggesting that it is dependent on FGF signaling (Fig. 4b). Given that FGF is not added to the medium during this period, ERK activation is most likely downstream of the FGF ligands FGF8 and FGF17, which are expressed by the cultures at these stages (Fig. 4c-d). We similarly observed nuclear localization of β-catenin on day 1 and 2, suggesting that Wnt signaling is mostly active during this period (Fig. 4a). Unexpectedly, nuclear β-catenin was also strongly down-regulated on day 3 despite the presence of the Wnt agonist Chir in the medium (Fig. 4a). These signaling dynamics suggest that differentiating iPS cells are exposed to transient Wnt/FGF signaling as in the posterior PSM *in vivo* (Fig. 4a), and that the regulation of FGF and Wnt signaling is largely autonomous in PSM cells differentiating *in vitro*. Thus, the arrest of *HES7-Achilles* oscillations observed on day 3 correlates with a strong reduction in FGF and Wnt signaling levels that likely brings about the onset of *MESP2* and *RIPPLY2* expression (Fig. 1i-j) as observed *in vivo^1^*. Indeed, treatment with the MEK1/2 inhibitor PD0325901 (PD03) for 24 hours caused most cells to decrease TBX6 expression and prematurely express *MESP2* (Fig. 4e and Extended Data Fig. 7a), indicating that FGF controls PSM maturation.

**Figure 4.**
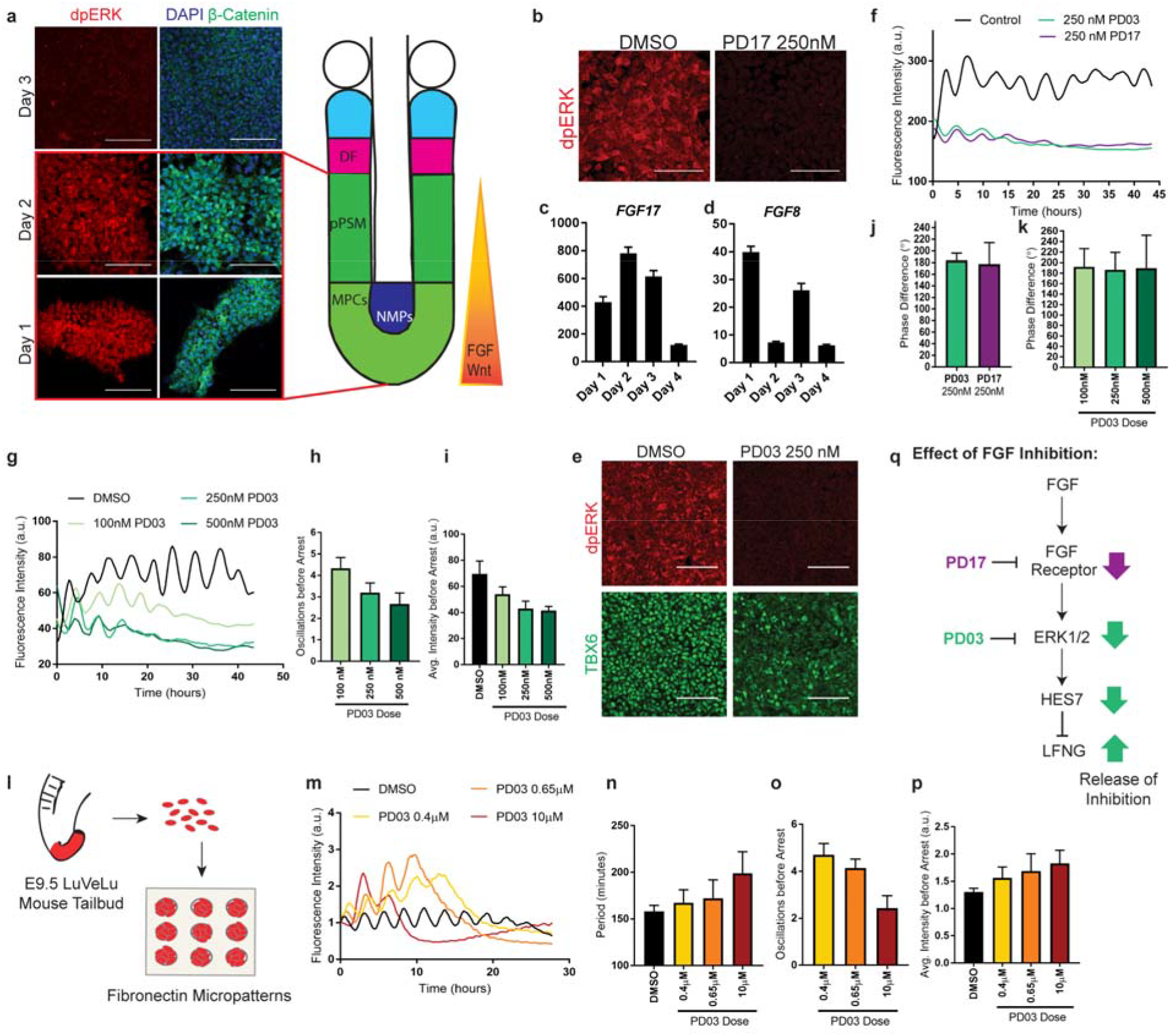
FGF signaling regulates the period and the phase of the segmentation clock. **a,** Left: Immunofluorescence staining for doubly phosphorylated ERK on days 1-3 of differentiation. Right: Immunofluorescence staining for β-catenin and nuclear staining on days 1-3 of differentiation. Nuclear localization of β-catenin is suggestive of active Wnt signaling. **b,** Immunofluorescence staining for doubly phosphorylated ERK on Day 2 of differentiation following 3 hours of treatment with either DMSO (vehicle control) or the FGFR inhibitor PD173074 (250 nM). **c,** qRT-PCR for the FGF ligand (*FGF17)* on days 1-4 of differentiation. Relative expression expressed as fold change relative to iPS (day 0). **d,** qRT-PCR for the FGF ligand (*FGF8)* on days 1-4 of differentiation. Relative expression expressed as fold change relative to iPS (day 0). **e,** Top: Immunofluorescence staining for doubly phosphorylated ERK on day 3 of differentiation (CLFBR medium) in cells treated with either DMSO or PD03 (250 nM). Bottom: Immunofluorescence staining for TBX6 on day 3 of differentiation (CLFBR medium) in cells treated with either DMSO or PD03 (250 nM). **f,** *HES7-Achilles* fluorescence intensity over the course of 45 hours in cells treated with DMSO (vehicle control), the MAPK inhibitor PD0325901 (250nM), or the FGFR inhibitor PD173074 (250nM) in CLFBR medium. **g,** *HES7-Achilles* fluorescence intensity profiles in cells treated with increasing doses of PD03 (100nM, 250 nM, 500nM) or vehicle control (DMSO). **h,** Average number of *HES7-Achilles* oscillations before arrest in cells treated with increasing doses of PD03 (100nM, 250 nM, 500nM). **i,** Average *HES7-Achilles* fluorescence intensity over the course of the oscillatory regime (i.e. prior to the arrest of oscillations) in cells treated with either vehicle control (DMSO) or increasing doses of PD03 (100nM, 250 nM, 500nM). **j,** Quantification of the average phase difference (degrees) for *HES7-Achilles* oscillations in PD03-or PD17-treated cells relative to control (DMSO) cells. **k,** Average phase difference (degrees) for *HES7-Achilles* oscillations in cells treated with increasing doses of PD03 (100nM, 250 nM, 500nM) relative to control (DMSO-treated) cells. **l,** Outline of the experimental strategy used to assess the effect of FGF inhibition in primary mouse PSM cells carrying the *LuVeLu* reporter. The tailbud is dissected from E9.5 transgenic embryos and cells are dissociated for seeding on fibronectin micropatterns. Oscillations of the *LuVeLu* reporter are examined in each micropattern. **m,** *LuVeLu* fluorescence intensity profiles in mouse tailbud explant cells cultured on CYTOO micropatterns in CLFBR medium containing DMSO (vehicle control) or increasing doses of PD03 (0.4μM, 0.65μM, 10μM). **n,** Average period of *LuVeLu* oscillations in mouse tailbud explant cells cultured on CYTOO micropatterns treated with DMSO (vehicle control) or increasing doses of PD03 (0.4μM, 0.65μM, 10μM).**o,** Average number of *LuveL*u oscillations before arrest in mouse tailbud explant cells cultured on CYTOO micropatterns treated with DMSO (vehicle control) or increasing doses of PD03 (0.4μM, 0.65μM, 10μM). **p,** Average *LuVeLu* fluorescence intensity over the course of the oscillatory regime (i.e. prior to the arrest of oscillations) in mouse tailbud explant cells cultured on CYTOO micropatterns treated with DMSO (vehicle control) or increasing doses of PD03 (0.4μM, 0.65μM, 10μM). **q,** Pathway illustrating the effect of FGF inhibition on the levels of FGFR and ERK activity, as well as overall levels of *HES7* and *LFNG*. Note that FGF inhibition by PD17 or PD03 leads to a decrease in *HES7* levels but an increase in *LFNG* levels due to release of inhibition. Mean ±SD; Scale bar = 100μm

In classical segmentation models such as the Clock-and-Wavefront, a maturation wave implemented by the FGF signaling gradient acts independently of the Clock to establish a specific threshold called Wavefront or determination front ^1,5^. At this level, FGF concentration becomes too low to sustain oscillations leading to their arrest thus defining the future segment ^1,2,5^. To directly test this hypothesis, we analyzed the effect of inhibiting FGF signaling on oscillations by treating *HES7-Achilles* reporter cells on day 2 with PD03 or PD17 in CLFBR medium. Unexpectedly, this did not result in an immediate arrest of oscillations but in their strong dampening followed by delayed arrest (Fig. 4f, Movie S7). This effect was dose-dependent with higher doses resulting in faster dampening and fewer oscillations before arrest (Fig. 4g-i). In addition, oscillations in treated cells exhibited a 180^0^ phase-shift relative to control cells resulting in treated cells and control cells to oscillate in antiphase, an effect which was not dose-dependent (Fig. 4j-k). To test whether this role of FGF is conserved in mouse, we used an *ex-vivo* system of PSM explants of the mouse reporter line *LuVeLu* that comprises a fluorescent construct reporting the oscillations of the cyclic gene *Lunatic fringe^4^*. Treating explants with 2 μM PD03 led to a transient increase in reporter intensity and delayed arrest of oscillations leading to the complete loss of *LuVeLu* signal (~ 4 oscillations were still observed after treatment; Extended Data Fig. 7b-d). To bypass the spatial effects inherent to this explant system, we used micropatterned cultures, where explant cells are dissociated and seeded on fibronectin disks (Fig. 4l). As with the human reporter cells, treating mouse cultures with FGF inhibitors led to a dose dependent effect on the number and intensity of oscillations (Fig. 4m-p). The period and average intensity of reporter oscillations progressively increased with inhibitor dose, a behavior reminiscent of *Lfng* oscillations during PSM maturation *in vivo* ^29^(Fig. 4m-p). In mouse, *Hes7* has been shown to repress *Lfng*^27^, thus providing a plausible explanation for the opposite response to FGF inhibition in terms of reporter intensity for the *HES7* and *Lfng* reporters (Fig. 4q). Thus, FGF signaling does not simply control the arrest of clock oscillations at the wavefront level as proposed in classical clock and wavefront models^1,5^. Overall, our data support the idea that FGF activity regulates the dynamics (period, phase and amplitude) of cyclic genes oscillations and the establishment of phase-gradients responsible for the traveling waves of the PSM in mammalian embryos.

Thus our work provides evidence for the existence of a human segmentation clock demonstrating the conservation of this oscillator from fish to human. We identify the human clock period as 5 hours indicating that the human segmentation clock operates roughly 2.5 times slower than its mouse counterpart^29^. This is consistent with the known difference in developmental timing between mouse and human embryos ^30^. Our culture conditions allow the production of an unlimited supply of cells displaying synchronized oscillations of the human segmentation clock and recapitulation of all major paraxial mesoderm stages. This therefore represents an ideal system to dissect the dynamical properties of the oscillator and its dysregulation in pathological segmentation defects such as congenital scoliosis.

**SUPPLEMENTARY INFORMATION** is available in the online version of the paper.

## ACKNOWLEDGMENTS

We thank members of the Pourquié lab, D. Ish-Horowicz, A. Klein and M. Heiman for critical reading of the manuscript and discussions. We thank Y. Saga for sharing reagents. Research in the Pourquié lab was funded by a grant from the National Institute of Health (5R01HD085121). D.E.W. is supported by 1K99GM121852.

## AUTHOR CONTRIBUTIONS

M.D.C. designed, performed and analyzed the biological experiments with O.P.; D.E.W. analyzed the single cell RNAseq data. C.B. optimized the dissociation protocol for single RNAseq and contributed to experiments with M.D.C. A.H. performed the mouse experiments. J. T. performed the mathematical analysis of synchronization. A.M. helped with quantifications. Z.A.T. generated the MSGN1-YFP line and helped M.D.C. with generation of the HES7-Achilles line. K.Y.K. and R.K. generated the destabilized Achilles construct. Y.N. and A.M. generated the Achilles fluorescent protein. M.D.C., D.E.W., A.H., C.B. and O.P. wrote the manuscript; O.P. supervised the project. All authors discussed and agreed on the results and commented on the manuscript.

## AUTHOR INFORMATION

Correspondence and requests for materials should be addressed to O.P.

## METHODS

### Generation of *HES7-Achilles* iPS reporter lines

The CRISPR/Cas9 system for genome editing was used to target the *HES7* locus in NCRM1 iPS cells^31^. A single guide RNA (Sense: 5’ ACCTGCTCGCCCGGACGCCC**GGG** 3’; protospacer adjacent motif (PAM) site highlighted in bold) targeting the 3’ end of *HES7* was designed using the MIT Crispr Design Tool (www.crispr.mit.edu) and cloned into the pSpCas9(BB)GFP vector (Addgene)^31^. This guide RNA was validated to efficiently cleave the target site with the T7 endonuclease 1 assay (T7E1; NEB cat. no. M0302S). T7E1 digestion of an 871 bp amplicon surrounding the target site (Fwd Primer 5’ GCTGCTACTTGTCCGGTTTCC 3’; Rev Primer 5’ TCGATCTCAGTTCCCGCTCTG 3’) revealed cleaved products roughly half the size of the intact fragment when genomic DNA was taken from NCRM1 iPS cells transiently expressing the pSpCas9(BB)GFP vector carrying the sgRNA. We also generated a repair vector consisting of 1kb 3’ and 5’ homology arms flanking a self-cleaving T2A peptide sequence, followed by the fast-folding YFP variant Achilles^16^, two destabilization domains (CL1 and PEST), and a nuclear localization signal (T2A-Achilles-NLS-CL1-PEST) in a pUC19 vector backbone by means of Gibson assembly (NEB). All primers used for Gibson assembly are listed in Extended Data Table 1. The assembled repair vector was then mutated by site directed mutagenesis to eliminate the PAM site (GGG>GGT) in the 3’ homology arm using the In-Fusion cloning kit (Takara). Both the pSpCas9(BB)GFP and targeting vector were delivered to iPS cells by nucleofection using a NEPA 21 electroporator. 24 hours after nucleofection, cells were sorted by GFP expression using an S3 cell sorter (Biorad) and seeded at low density in Matrigel-coated plates (Corning, cat. no. 35277) in mTeSR1 (Stemcell Technologies cat. no. 05851) + 10μM Y-27362 dihydrochloride (Tocris Bioscience, cat. no. 1254). Single cells were allowed to expand clonally and individual colonies were screened by PCR for targeted homozygous insertion of 2A-Achilles-CL1-PEST-NLS immediately before the stop codon of *HES7*. The following primers anneal outside the repair vector homology arms and were used for genotyping: Fwd 5’ ATCTCCTCCTCACGCGTTGG 3’; Rev 5’ AGAGTGCCAAATTGATTCGTCTCC 3’. Positive clones were sequenced to ensure no undesired mutations in the *HES7* locus had been introduced by the genome editing process. Three homozygous clones were further validated by RT-qPCR and immunofluorescence.

### iPS cell culture and 2D differentiation

NCRM1 iPS cells (RUCDR, Rutgers University) and lines carrying the *MSGN1-Venus*^13^ or *HES7-Achilles* reporters were maintained in Matrigel-coated plates (Corning, cat. no. 35277) in mTeSR1 medium (Stemcell Technologies cat. no. 05851) as previously described^10^. Paraxial mesoderm differentiation was carried out as described^10^. Briefly, mature iPS cell cultures were dissociated in Accutase (Corning cat. no. 25058CI) and seeded at a density of 3 × 10^4^ cells per cm^2^ on Matrigel-coated plates in mTeSR1 and 10μM Y-27362 dihydrochloride (Rocki; Tocris Bioscience, cat. no. 1254). Cells were cultured for 24-48 hours until small, compact colonies were formed. Differentiation was initiated by switching to CL medium consisting of DMEM/F12 GlutaMAX (Gibco cat. no. 10565042) supplemented with 1% Insulin-Transferrin-Selenium (Gibco cat. no. 41400045), 3 μM Chir 99021 (Tocris cat. no. 4423) and 0.5 μM LDN193189 (Stemgent cat. no. 04-0074). On day 3 of differentiation, cells were changed to CLF medium consisting of CL medium with 20ng/ml murine bFGF (PeproTech cat. no. 450-33). Media was changed daily.

For live imaging experiments, differentiation was performed as described above, except cells were seeded on 35 mm matrigel-coated glass-bottom dishes (MatTek cat. no. P35G-1.5-20-C) or 24 well glass-bottom plates (In Vitro Scientific cat. no. P24-1.5H-N). DMEM/F12 without phenol red was used to reduce background fluorescence (Gibco cat. no. 21041025).

To extend the oscillatory window of differentiated PSM cells, we cultured *HES7-Achilles* cells in CLFBR medium consisting of DMEM/F12 GlutaMAX, 1% ITS, 3 μM Chir 99021, 0.5 μM LDN193189, 50 ng/ml mFgf4 (R&D Systems cat. no. 5846-F4-025), 1 μg/ml Heparin (Sigma Aldrich cat. no. H3393-100KU), 2.5 μM BMS493 (Sigma Aldrich cat. no. B6688-5MG) and 10 μM Y-27362 dihydrochloride starting on day 2 of differentiation. Media was refreshed daily.

To track oscillations in individual cells within the culture, we mixed *HES7-Achilles* cells with NCRM1 cells with a ratio of 1:100 at the time of seeding for pre-differentiation. Cells were then differentiated normally under CLFBR conditions.

To examine oscillations in isolated cells, we differentiated *HES7-Achilles* cells normally (CL medium) for the first two days on 35mm plastic dishes and dissociated them with Accutase (Corning cat. no. 25058CI) on day 2 of the differentiation protocol. Cells were reseeded on fibronectin-coated (BD Biosciences cat. no. 356008) or bovine serum albumin (BSA)-coated (Gibco cat. no. 15260-037) 24 well glass-bottom plates at high (500,000 cells per well) or low density (25,000-50,000 cells per well) in CLFBR media. Using our regular DMEM/F12 base media resulted in poor survival of low density cultures. We found that using RHB Basal media (Takara/Clontech cat. no. Y40000), supplemented with 5% knockout serum replacement (KSR; Thermo Fisher cat. no. 10828-028) improved survival significantly.

### Explant culture

Explant culture was performed as previously described^4^. *LuVeLu* E9.5 mice were sacrificed according to local regulations in agreement with national and international guidelines. Study protocol was approved by Brigham and Women’s Hospital IACUC/CCM/ Ectoderm was removed using Accutase (Life Technologies) and tailbud was dissected with a tungsten needle. Explants were then cultured on fibronectin-coated plate (LabTek chamber). The medium consists of DMEM, 4.5g/L Glucose, 2mM L-Glutamine, non-essential amino acids 1x (Life Technologies), Penicillin 100U/mL, Streptomycin 100μg/mL, 15% fetal bovine serum (FBS), Chir-99021 3μM, LDN193189 200nM, BMS-493 2.5 μM, mFgf4 50ng/mL, heparin 1μg/mL, HEPES 10mM and Y-27632 10μM. Explants were incubated at 37°C, 7.5% CO2. Live imaging was performed on a confocal microscope Zeiss LSM 780, using a 20X objective (note that the tiling could create lines between the different images). For micropattern culture, explants were cultured overnight in standard condition, then dissociated using Trypsin-EDTA, and plated on fibronectin-coated CYTOOchips Arena in a CYTOOchamber 4 wells.

### Small Molecule Inhibitor Treatments

To inhibit Notch signaling, 25 μM DAPT (Sigma Aldrich cat. no. D5942-5MG) was added to CLFBR media on day 2 of differentiation. To inhibit FGF signaling, PD0325901 (Stemgent 04-006) or PD173074 (Cayman Chemical cat. no. 219580-11-7) were added to CL or CLFBR media at the indicated concentrations. Latrunculin A (Cayman Chemical ca. no. 10010630), which inhibits actin polymerization and YAP signaling, was used at 350 nM in RHB basal media supplemented with CLFBR and 5% KSR. Mouse explants and micropatterned cultures were treated with PD0325901 (Sigma-concentration as described in the text) and PD173074 (Sigma - 250nM).

### Time-lapse Microscopy

Time lapse-imaging of *MSGN1-Venus* or *HES7-Achilles* PSM cells was performed on a Zeiss LSM 780 point-scanning confocal inverted microscope fitted with a large temperature incubation chamber and a CO_2_ module. An Argon laser at 514 nm and 7.5% power was used to excite the samples through a 20X Plan Apo (N.A. 0.8) objective. Images were acquired with an interval of 18 minutes for a total of 24-48 hours. A 3×3 tile of 800×800 pixels per tile with a single z-slice of 18 μm thickness was acquired per position. Multiple positions, with at least two positions per sample, were imaged simultaneously using a motorized stage. Explant imaging was performed on a Zeiss LSM780 microscope using a 20X/0.8 objective. For mouse cells imaging, single section (~19.6μm wide) with tiling (3×3) of a 512×512 pixels field was acquired every 7.5 minutes (in most experiments) at 8-bit resolution.

### Immunostaining

For immunostaining of 2D cultures, cells were grown on Matrigel-coated glass-bottom plates or 12mm glass coverslips placed inside plastic dishes. Cells were rinsed in Dulbecco’s phosphate buffered saline (DPBS) and fixed in a 4% paraformaldehyde solution (Electron Microscopy Sciences cat. no. 15710) for 20 minutes at room temperature, then washed 3 times with phosphate buffered saline (PBS). Typically, samples were permeabilized by washing three times for three minutes each in Tris buffered saline (TBS) with 0.1% Tween (known as TBST) and blocked for one hour at room temperature in TBS-0.1% Triton-3% FBS. Primary antibodies were diluted in blocking solution and incubated overnight at 4°C with gentle rocking. Primary antibodies and dilution factors are listed in Ext. Data Table 2. Following three TBST washes and a short 10-minute block, cells were incubated with Alexa-Fluor conjugated secondary antibodies (1:500) and Hoechst33342 (1:1000) overnight at 4°C with gentle rocking. Three final TBST washes and a PBS rinse were performed, and cells were mounted in Fluoromount G (Southern Biotech cat. no. 0100-01). Images were acquired using either a Zeiss LSM880 or LSM780 point scanning confocal microscope with a 20X objective.

For visualizing phospho-ERK1/2 in 2D monolayer differentiated cells, cells were transferred onto ice and quickly rinsed in ice-cold PBS containing 1 mM sodium vanadate (NaVO_4_). Next, cells were fixed in 4% paraformaldehyde for 15 min at room temperature, rinsed three times in PBS and dehydrated in cold methanol at −20°C for 10 minutes. Following three PBS rinses, cells were blocked in PBS containing 0.1% Triton X-100 and 5% goat serum and incubated in pERK1/2 antibody diluted in antibody buffer (0.1% Triton X-100 and 1% BSA in PBS) overnight at 4°C. Cells were washed in PBS, and incubated in blocking solution for 10 minutes and with secondary antibody and Hoechst33342 in antibody buffer overnight at 4°C. Cells were rinsed three times in PBS before mounting and imaging as described above.

### RNA extraction, reverse transcription and qPCR

Cells were harvested in Trizol (Life Technologies cat. no. 15596-018), followed by precipitation with Chloroform and Ethanol and transfer onto Purelink RNA Micro Kit columns (Thermo Fisher cat. no. 12183016) according to manufacturer’s protocol, including on-column DNase treatment. A volume of 22 μl RNase-free water was used for elution and RNA concentration and quality were assessed with a Nanodrop. Typically, between 0.2-1 μg of RNA was reverse transcribed using Superscript III First Strand Synthesis kit (Life Technologies cat. no. 18080-051) and oligo-dT primers to generate cDNA libraries.

For real time quantitative PCR, cDNA was diluted 1:30-1:50 in water and qPCR was performed using the iTaq Universal SYBR Green kit (Bio-Rad cat. no. 1725124). Each gene-specific primer and sample mix was run in triple replicates. Each 10 μl reaction contained 5 μl 2X SYBR Green Master Mix, 0.4 μl of 10 μM primer stock (1:1 mix of forward and reverse primers), and 4.6 μl of diluted cDNA. qPCR plates were run on a Bio-Rad CFX384 thermocycler with the following cycling parameters: initial denaturation step (95°C for 1 minute), 40 cycles of amplification and SYBR green signal detection (denaturation at 95°C for 5 seconds, annealing/extension and plate read at 60°C for 40 seconds), followed by final rounds of gradient annealing from 65°C to 95°C to generate dissociation curves. Primer sequences are listed in Extended Data Table 3. All unpublished primers were validated by checking for specificity (single peak in melting curve) and linearity of amplification (serially diluted cDNA samples). For relative gene expression analysis, the ΔΔCt method was implemented with the CFX Manager software. *PP1A* was used as the housekeeping gene in all cases. Target gene expression is expressed as fold change relative to undifferentiated iPS cells.

### Flow Cytometry Analysis

To determine the fraction of PSM cells expressing *MSGN1-Venus*, cultures were dissociated in Accutase and analyzed by flow cytometry using an S3 cell sorter (Biorad). Undifferentiated *MSGN1-Venus* iPS cells, which do not express the fluorescent protein, were used as a negative control for gating purposes. Samples were analyzed in biological triplicates. Results are presented as the percentage of Venus positive cells in the sorted fraction.

### Image Analysis

Time lapse movies of *HES7-Achilles* were first stitched and separated into subsets by position in Zen. Then, background subtraction and Gaussian blur filtering were performed in ImageJ to enhance image quality. A small region of interest (ROI) was drawn and the mean fluorescence intensity over time was calculated. Intensity is presented in arbitrary units, which are sometimes regularized to oscillate around zero by subtracting the moving average with a window size of 10 timepoints. For smoothening, we applied the Sgolay filtering algorithm in MATLAB. Following moving average subtraction, we performed Fourier transformation of *HES7-Achilles* intensity profiles to determine the predominant period of oscillations. The Hilbert transformation was used to calculate the instantaneous frequency and phase of *HES7-Achilles* oscillations. To compare the phase between DMSO and PD17 or PD03 treated cells, we used the Hilbert transformation to calculate the instantaneous phase of each curve separately, then subtracted the phase of treated cells from untreated cells at each timepoint. Phase difference is expressed as the average of instantaneous phase differences before the arrest of oscillations in treated cells. To track oscillations in isolated or sparse *HES7-Achilles* cells in a NCRM1 background, we manually tracked cells by drawing a circle around the nucleus of an individual cell at each time point and measuring fluorescence intensity inside the ROI.

For mouse explants, kymographs were done in Fiji by drawing a rectangle from the starting center of the traveling waves to the edge of the explant perpendicular to the direction of the wave. The intensity along the long axis was measured and the image was smoothened (this filter replaces each pixel with the average of its 3 × 3 neighborhood).

Fluorescence intensity profiles were done by selecting a circular region of interest in FiJi and by measuring the total intensity over time for this region; *LuVeLu* intensity is given in arbitrary units (normalized by the initial value) and a smoothing function (average over three points) was applied. Fluorescence intensity shows the mean fluorescence smoothed by applying a moving average over five points (with equal weight). For the quantification of micropattern experiments, a region of interest encompassing the entire surface of one circle was drawn and the *LuVeLu* intensity was measured using the Time Series Analyzer V3 plugin on Fiji. The period was measured by measuring the time between two peaks or two troughs. The average intensity was measured by averaging the intensity over 3 hours to avoid instantaneous variations dues to the oscillations.

### Phase and phase shifts

Synchronization was evaluated from the fluorescence intensity data. The data show clear oscillations with relatively heterogeneous profiles and background fluorescence intensity (see Movie S5). To evaluate precisely an instantaneous phase, we used classical methods and developed custom algorithms adapted to the signals observed. All algorithms provided consistent results.

#### Method 1

we developed a local cross-correlation algorithm that computes the phase shift between two signals at a given time as the optimal timeshift of a local part of the signal within a time-window of 6 hours, slightly larger than the period of the average signal (the average distance between two peaks of the mean of all fluorescence trajectories yields a period of 5.02 hours). This algorithm was developed in Matlab. We used this cross-correlation algorithm to compute phase differences between individual signals and the global oscillatory pattern, either computing the phase shift between each pair of cells, or between one cell and the mean signal.

#### Method 2

Alternatively, we defined intrinsic phases for each signal as the relative time between two peaks. Peaks were detected using the *findpeaks* function of Matlab. When peak times are detected, say at times t_0_,t_1_,…,t_n_ the phase of the signal at a given time t ∈ [t_i_,t_i+1_] was defined as 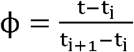. The code was developed in Matlab.

#### Method 3

We compared these two new methods, well adapted to the type of signal obtained, to the classical Hilbert transform method. This efficient method presents a few well-known artifacts, particularly due to changes in the shape of the signal, classically corrected using a detrending algorithm. To improve the evaluation of the phase using the Hilbert transform, we used a local detrending consisting in removing a moving average computed over a time window of 6h, allowing correcting for local trends, and evaluated the phase using the *hilbert* function of Matlab.

### Synchronization and the Kuramoto model

To quantify the level of synchrony of a given set of signals, we compared the distribution of phase shifts with the classical Kuramoto model^27^. This parameters: the number of interacting oscillators n, the disorder of intrinsic oscillation frequency model describes the dynamics of heterogeneous coupled oscillators, and depends on three ω and the coupling parameter K. Denoting θ_i_ the phase of oscillator i ∈ {1⋯n}and ω_i_ its frequency, the Kuramoto model describes the phases as:

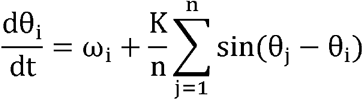

and the model classically shows two phases: synchrony when the coupling K is sufficiently strong respective to the disorder ω, or disorder. The synchronization of the system is well-described by the Kuramoto order parameter:

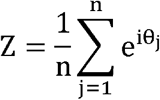

where i is the complex variable. Its argument provides the average phase of the system, while its modulus quantifies the level of synchrony: it is equal to 1 when all oscillators have the same phase, and 0 when the oscillators are evenly distributed.

Using the phases derived, we evaluated as a function of time the order parameter and its modulus. We compared this value to simulations of the original Kuramoto model and varied the parameters to find those values fitting closely the experimentally computed order parameter and its fluctuations.

For the control case, we found a good consistency with ω = 3.1, n = 500 and K = 1.12 K_c_ with 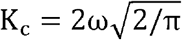 the critical coupling coefficient when the distribution of phases is symmetric. The initial condition was taken with a standard deviation of 0.14 radians.

For the DAPT condition, we found that the experimentally computed Kuramoto order parameter was consistent with ω = 1.8, n = 50 and K = 0.8 K_c_, and an initial condition with a standard deviation of 0.125 radians.

### Quantification of cell diffusivity

The diffusivity was measured by manually tracking cells. The two-dimensional mean squared displacement 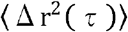 where 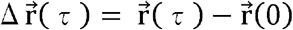 is the displacement after a lag time τ, was computed using the overlapping intervals method ^32^. We verified that the trajectories were following a random-walk law, as the mean square displacement exhibits a linear trend with respect to the lag time. We therefore extracted the diffusivity D by linear regression:

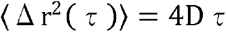

### Preparation of single-cell suspensions for single cell RNA-sequencing

Single-cell dissociation protocols for the various tissues and cells analyzed were optimized to achieve >90% viability and minimize doublets prior to sample collection.

For hiPS differentiation, 5×10^4^ *MSGN1-Venus* cells were seeded on Matrigel-coated 24-well plates 48 hours prior to differentiation. Cells were differentiated according to our previously published protocol^10^. All samples (Day 1-4 and hiPS control samples) were dissociated, collected and captured on an inDrops setup on the same day, two biological replicates per sample. For dissociation, cells were briefly rinsed in PBS, and incubated in TrypLE Express (Gibco) for 5 min at 37°C. Dissociated cells were run through a 30 μm cell strainer, spun down at 200g for 4 min at 4°C and resuspended in 100 μl 0.5% BSA in PBS.

For generating cell suspensions from mouse embryo tailbuds, E9.5 embryos (25-28 somite stage) from CD-1 IGS mice (Charles River) were collected and the posterior part of the embryo, including the last 3 pairs of somites, was carefully dissected from two littermate embryos and subsequently processed as separate samples. Tissues were collected in PBS and dissociated in TrypLE Express for 10 min at 37°C. Cells were rinsed in PBS/EDTA, transferred to 0.5% BSA in PBS, mechanically separated by trituration and run through a 30 μm cell strainer. Cells were spun down at 200 g for 4 min at 4°C and resuspended in 100 μl 0.5% BSA in PBS.

The following number of cells were sequenced per sample:

1. 2D differentiation samples (two biological replicates processed independently) For each replicate hiPS control, 1000 cells; Day 1, 1500 cells; Day 2, 1500 cells; Day 3, 1500 cells; Day 4, 1500 cells
2. Embryo samples tailbud cells from two E9.5 embryos, (2x 3000 cells processed independently) Every sample was collected as biological replicate and sequencing data from both samples were combined for data analysis. Actual number of cells captured on inDrops was twice as many as sequenced for backup purposes.

### Barcoding, sequencing, and mapping of single-cell transcriptomes

Single-cell transcriptomes were barcoded using inDrops ^20^ as previously reported ^33^, using “V3” sequencing adapters. Following within-droplet reverse transcription, emulsions consisting of ~1,000-3,500 cells were broken, frozen at −80C, and prepared as individual RNA-seq libraries. inDrops libraries were sequenced on an Illumina NextSeq 500 using the NextSeq 75 High Output Kits using standard Illumina sequencing primers and 61 cycles for Read1, 14 cycles for Read2, 8 cycles each for IndexRead1 and IndexRead2. Raw sequencing data (i.e. FASTQ files) were processed using the inDrops.py bioinformatics pipeline available at github.com/indrops/indrops. Transcriptome libraries were mapped to human or mouse reference transcriptomes built from the GRCh37 / hg19 (GCF_000001405.13) or GRCm38 / mm10 (GCF_000001635.20) genome assemblies, respectively. Bowtie version 1.1.1 was used with parameter –e 200.

### Processing of single-cell RNA-seq data

inDrops data were filtered to only include transcript counts originating from abundant cell barcodes. This determination was performed by inspecting a weighted histogram of Unique Molecular Identifier (UMI) – gene pair counts for each cell barcode, and manually thresholding to include the largest mode of the distribution (in all cases >80% of total sequencing reads). Transcript UMI counts for each biological sample were then reported as a transcripts x cells table and adjusted by a total-count normalization. For each UMI counts table, highly variable genes were identified as follows: (1) fano factors were calculated for all genes; (2) genes were then ranked by an above-Poisson noise statistic^20^; (3) the top 2000 variable genes according to this statistic were filtered to exclude genes whose single-cell transcript counts were only weakly correlated (correlation coefficient < 0.2) to any other variable gene. Sets of cell cycle and housekeeping-associated genes were excluded from downstream analyses. These sets were generated by expanding a list to include any gene whose expression was highly correlated (correlation coefficient > 0.4) to any of the following genes. Mouse: Rpl5, Rps6, Rps8, Rplp1, Mcm6, Mcm7, Plk1, Cdk1, Aurka, Cenpa, Cenpf, Mcm4, Rrm2, Hmgb1, Hmga2, Cdc6, Hspa8, Hsp90ab1, Hspe1; Human: RPSA, RPS5, RPS9, RPS17, RPS18, RPSA, RPS3A, RPL31, RPL7A, RPS25, RPL3, CDK1, CENPA, AURKA, PLK1, CENPF, TOP2A, MCM2, MCM3, MCM5, MCM6, RRM2, MCM7, HMGB1, EIF5, HSPE, BUB1, CEP55, KIF14, CDKN3, CDCA2, CCNF. The resulting lists were then subjected to a second “expansion” round, and all associated genes were discarded. Mouse E9.5 data were filtered for doublet-like cells by simulating synthetic doublets from pairs of scRNA-seq profiles, and assigning scores based on a k-nearest-neighbor classifier on the PCA-transformed data (see Extended Data Fig 2a-b).

### Low Dimensional Embedding and Clustering

Single-cell data were projected into a low dimensional subspace by first standardizing (by a z-score) the normalized UMI counts for each gene, followed by principal component analysis (PCA). The top significant PCA dimensions were estimated by comparing the distribution of eigenvalues to that of a randomized dataset, as previously described^20^; only significant principal components were analyzed. For Figs 2a, Extended Data Fig. 2a, and Extended Data Fig. 2c, single-cell RNAseq data were visualized by two-dimensional t-distributed stochastic neighbor embedding (tSNE)^33^ on cell PC scores using a perplexity setting of 30, with 1000 iterations. For Figs 2b-g, Extended Data Fig. 3, k-nearest-neighbor (knn) graphs were constructed from cell PC scores using a small variation of the SPRING method^33^, in which nearest neighbors were identified using correlation distance. For the human graphs, only mutual edges were retained. All kNN graphs were visualized in 2D using a force-directed layout^34^. For Figs 2d, 2e, Extended Data Fig. 3a, and Extended Data Fig. 3d, cell clusters were identified by GenLouvain, a graph-based community detection algorithm^35^. For tSNE maps appearing in Fig. 2a and Extended Data Fig. 2c, cell clusters were identified by density clustering^36^.

### Identification of Differentially Expressed Genes

Cluster-defining transcripts were identified by a Wilcoxon rank-sum test by comparing cells of each cluster to cells from all other clusters in the same sample. Genes were considered differentially expressed if they met the following criteria: absolute log2 fold-change >1, adjusted p-value < 0.05. P-values were adjusted for multiple hypotheses as previously reported ^37^. All tables report top-ranking differentially expressed genes, ranked by p-value.

### Pseudo-Spatiotemporal Ordering and Identification of Dynamically Varying Genes

A pseudo-spatiotemporal ordering of cells along the paraxial mesoderm (PM) lineage was constructed from the E9.5 nearest neighbor graph in a variation of Wanderlust^37^. First, a series of shortest paths were determined between cells of the NMP and Somite clusters; paths were restricted to a subgraph containing cells of the MPC, aPSM, and pPSM clusters in which 80% of all edges were randomly deleted. This process was repeated for a total of 100 iterations in which different sets of edges were deleted. All cells/nodes discovered during this process were ordered based on their average position over all shortest paths in which they appeared. Genes that varied dynamically along this trajectory were then identified as previously described^38^. Sliding windows of 100 cells were first scanned to identify two windows with maximum and minimum average expression levels for all genes, respectively. A t-test was then performed between these two sets of 100 expression measurements (FDR < 0.01). Gaussian-smoothened expression z-scores for significantly variable genes along the trajectory were then calculated; genes were ranked by peak expression and plotted as a heatmap. A subset of PM differentiation markers were highlighted on the heatmap. The full list of dynamically expressed genes appears in Table S2.

### Data Availability

Raw sequencing data, raw and normalized counts data, and single-cell clustering assignments are available from NCBI Gene Expression Omnibus (GEO), Accession # GSE114186 (Reviewer Token: ctgfocsavjgllcv). Interactive versions of the analyzed datasets can be accessed as follows.

Mouse E9.5 tSNE clustering analysis (Fig. 2a and Extended Fig. 2c): https://kleintools.hms.harvard.edu/tools/springViewer_1_6_dev.html?datasets/Pourquie2018/E95/full

Mouse E9.5 k-nearest neighbor graph of paraxial mesoderm and neural clusters (Fig 2b-f and Extended Fig 3a): https://kleintools.hms.harvard.edu/tools/springViewer_1_6_dev.html?datasets/Pourquie2018/E95_sm2/full

Human iPS cultures Day 0 – Day 4 (Fig 2e-g and Extended Figs. 3d, 4a-b) https://kleintools.hms.harvard.edu/tools/springViewer_1_6_dev.html?datasets/Pourquie2018/HS2D/full

### Code Availability

Single-cell sequencing data were processed and analyzed using publicly available software packages: https://github.com/indrops/indrops and https://github.com/AllonKleinLab/SPRING. Data processing and analysis routines are described in the methods. Matlab code for reproducing panels appearing in Fig 2 and Extended Figs 2-4 will be made available upon publication.

## EXTENDED DATA TABLES

**Extended Data Table 1.**
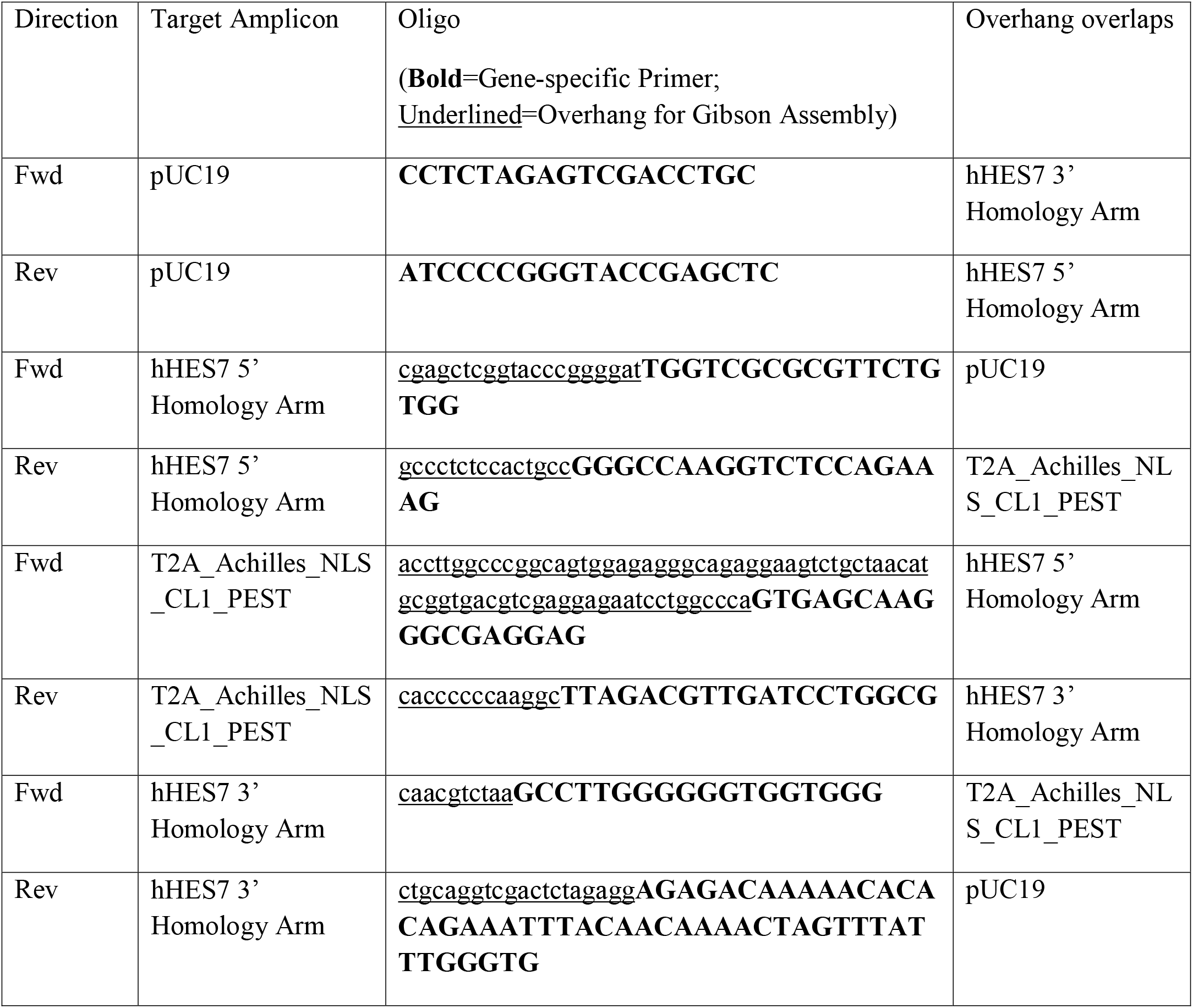
Primers used for Gibson assembly of *HES7-Achilles* repair vector

**Extended Data Table 2.** Primary Antibodies for Immunofluorescence

| Antibody | Species | Type | Source | Catalog Number | Dilution |
| --- | --- | --- | --- | --- | --- |
| OCT3/4 | Mouse | Monoclonal | Santa Cruz | Sc-5279 | 1:800 |
| SOX2 | Rabbit | Polyclonal | Millipore | AB5603 | 1:300 |
| T/BRACHYURY | Goat | Polyclonal | R&D | AF2085 | 1:300 |
| TBX6 | Rabbit | Polyclonal | Abcam | ab38883 | 1:300 |
| CDH1 | Mouse | Monoclonal | Abcam | ab76055 | 1:300 |
| CDH2 | Rabbit | Polyclonal | Abcam | ab12221 | 1:300 |
| dpERK | Rabbit | Monoclonal | Cell Signaling | 4370P | 1:400 |
| β-CATENIN | Mouse | Monoclonal | BD | 610153 | 1:400 |
| YAP | Mouse | Monoclonal | Santa Cruz | sc-101199 | 1:200 |

**Extended Data Table 3.**
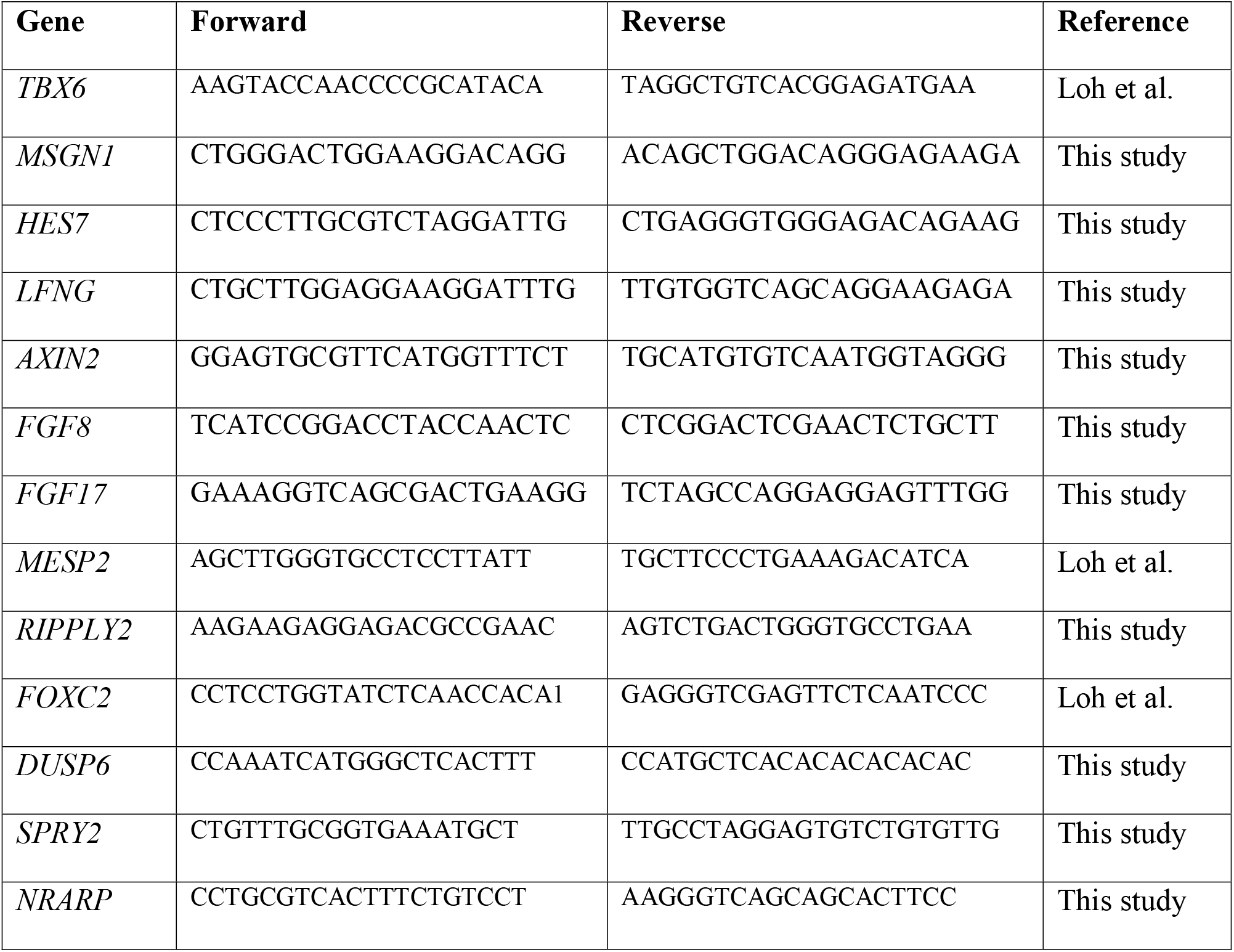
Primer sequences for qPCR

**Extended Data Figure 1.**
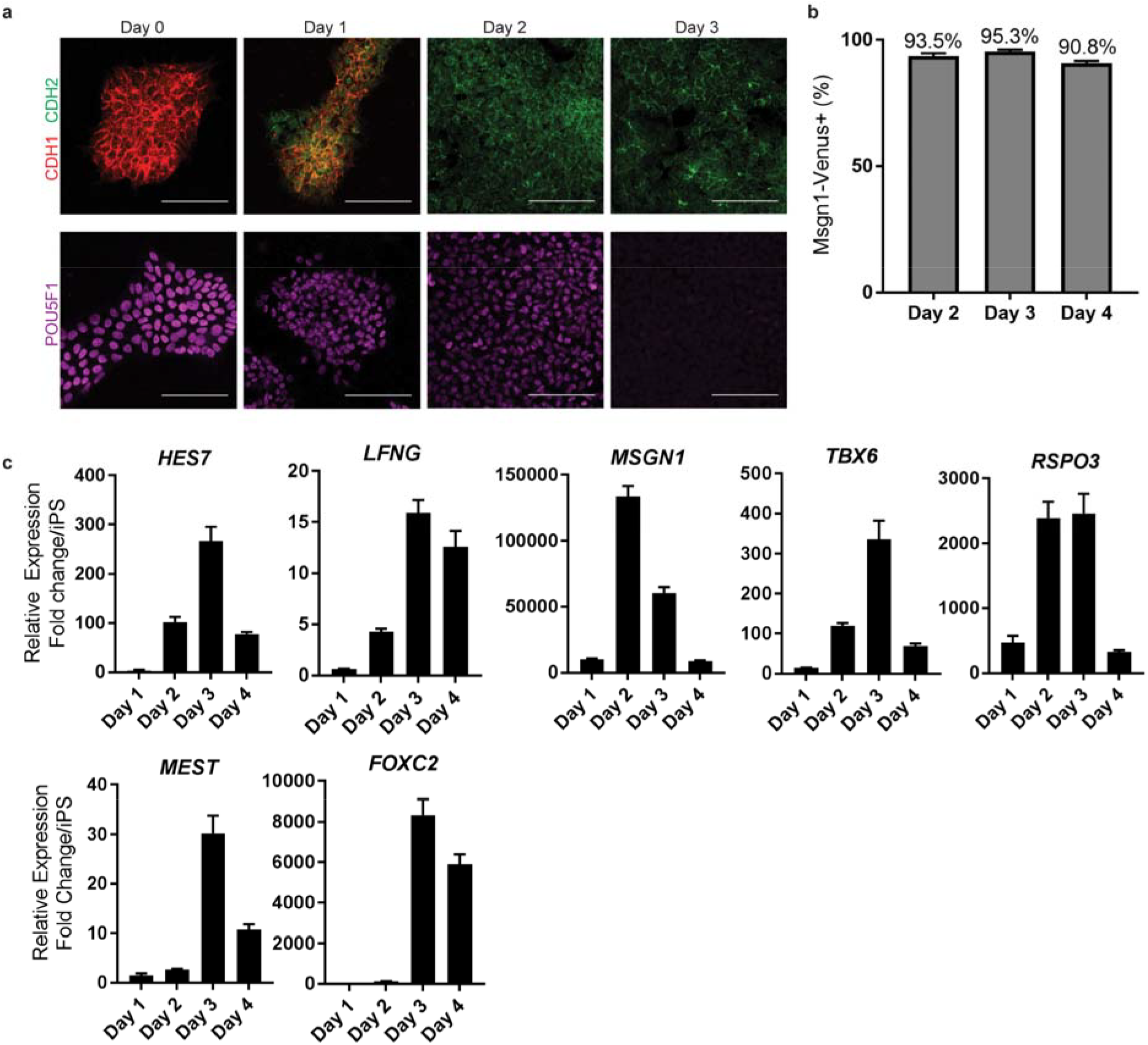
**a,** Top: Immunofluorescence staining for the cadherins CDH1 and CDH2 on days 0-4 of differentiation. Bottom: Immunofluorescence staining for the pluripotency factor POU5F1 on days 0-4 of differentiation. Scale bar = 100 μm**. b,** Percentage induction of the *MSGN1-Venus* reporter as determined by FACS on days 2-4 of differentiation. **c,** qRT-PCR for cyclic genes (*HES7, LFNG*), posterior PSM markers (*MSGN1, TBX6, RSPO3*), and anterior PSM markers (*MEST, FOXC2*) on days 1-4 of differentiation. Relative expression expressed as fold change relative to iPS (day 0). Mean ±SD.

**Extended Data Figure 2.**
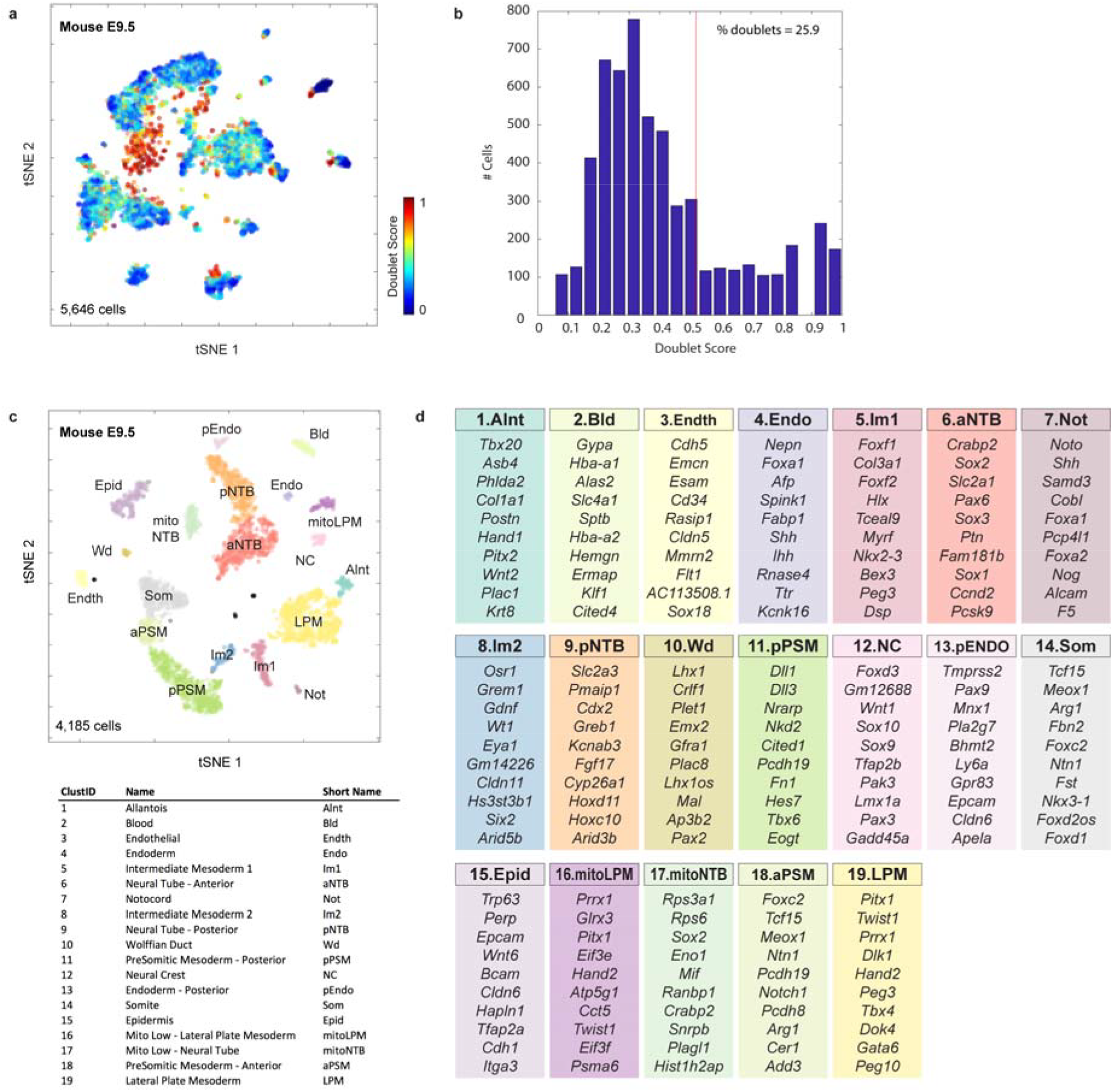
**a,** Pre-filtering of doublet-like cells. tSNE embedding shows all original E9.5 cells, colored by doublet score. Doublet scores indicate the extent to which a given single-cell transcriptome resembles a linear combination of two randomly selected cells (see Methods). **b,** Histogram of doublet scores. Scores >0.5 were filtered from subsequent analyses. **c,** tSNE embedding of E9.5 cells, post-doublet filtering. Individual cells are colored according to tSNE density cluster ID. **d,** Top 10 significantly enriched transcripts for each tSNE cluster. Reported transcripts are filtered (log2-fold change >1, adj. p-value < 0.05), and ranked by p-value (see also Table S1).

**Extended Data Figure 3.**
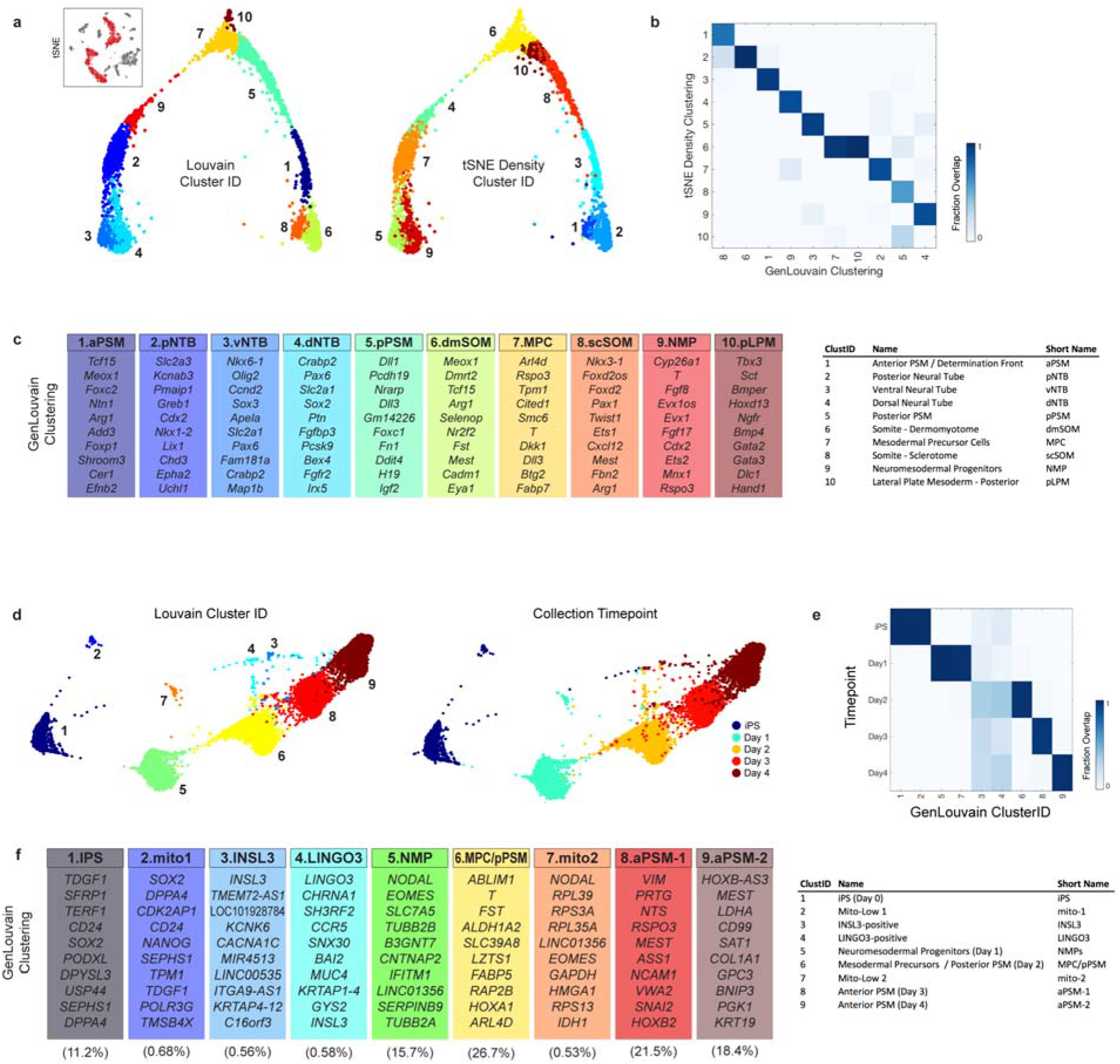
**a,** Mouse E9.5 kNN graph, colored by Louvain or tSNE/Density cluster IDs. **b,** Confusion matrix plots overlap of clustering assignments, normalized by column. **c,** Top 10 significantly enriched transcripts for each Louvain cluster. Reported transcripts are filtered (log2-fold change >1, adj. p-value < 0.05), and ranked by p-value. See also Table S3. **d,** Human iPS kNN graph, colored by Louvain cluster ID or collection timepoint. **e,** Confusion matrix (column normalized) plots overlap of timepoints and clusters. Excluding clusters 3-4, all other clusters map to a single timepoint. **f,** Top 10 significantly enriched transcripts for each Louvain cluster. Reported transcripts are filtered (log2-fold change >1, adj. p-value < 0.05), and ranked by p-value. See also Table S4.

**Extended Data Figure 4.**
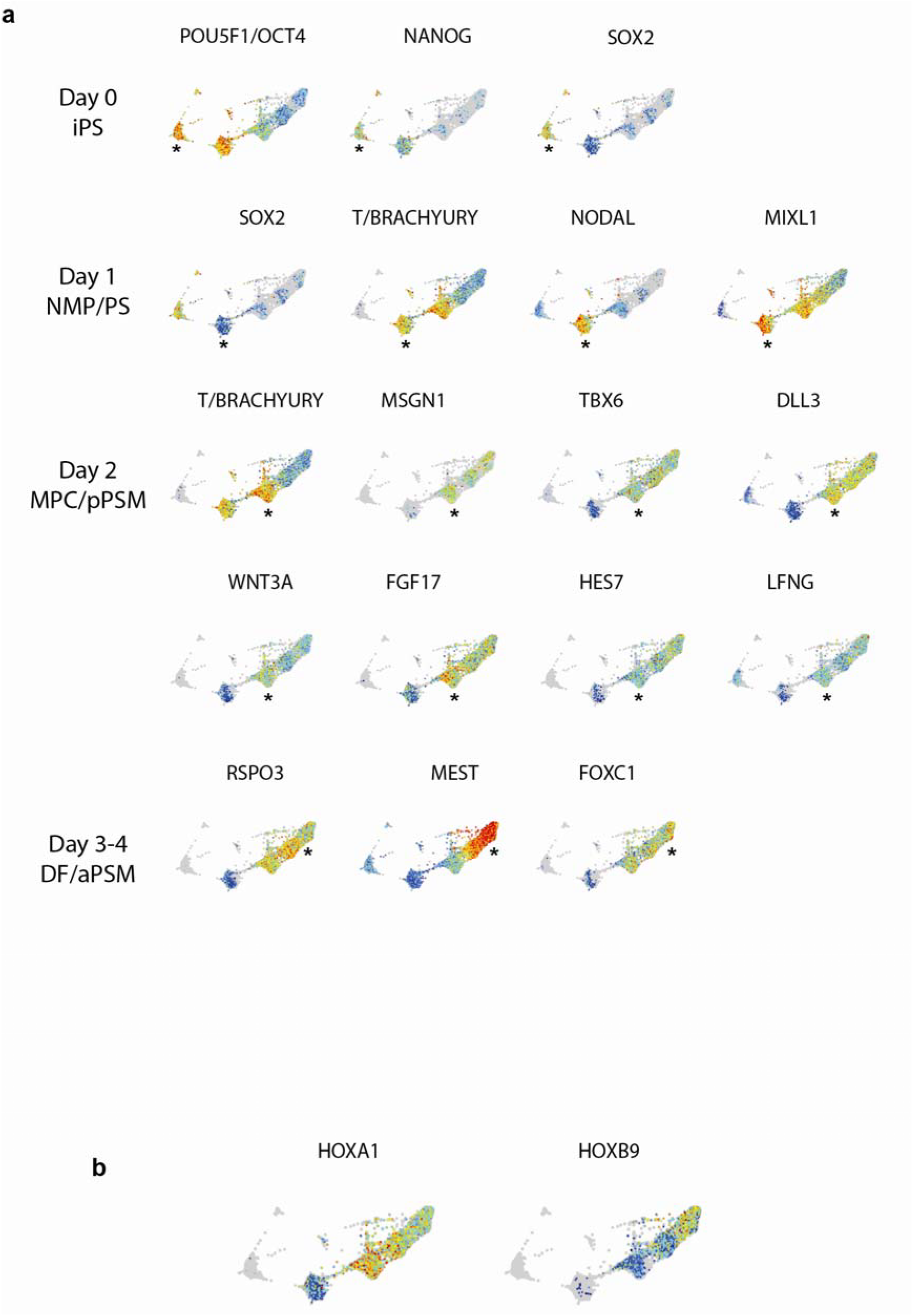
**a,** Expression patterns of signature genes during human iPS differentiation into PSM *in vitro*. Asterisks mark the cluster corresponding to the differentiation day indicated on the left. **b,** Expression pattern of Hox genes showing the delayed activation of *HOXB9* compared to *HOXA1.*

**Extended Data Figure 5.**
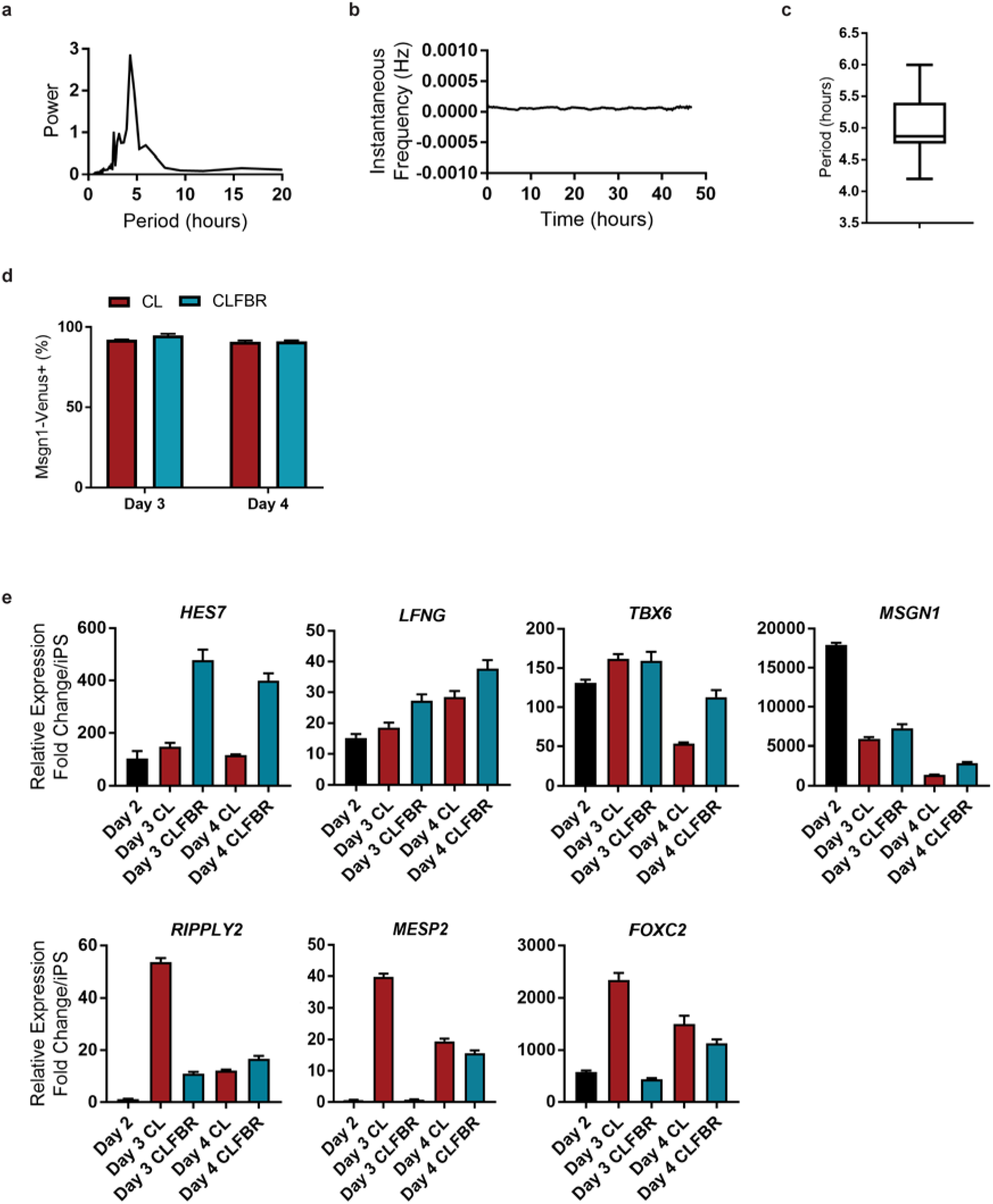
**a,** Fourier transform of *HES7-Achilles* oscillations under CLFBR conditions, indicating the major period of oscillations. **b,** Instantaneous frequency (Hz) over time for *HES7-Achilles* oscillations in CLFBR medium. **c,** Average period of *HES7-Achilles* oscillations in CLFBR medium. **d,** Comparison of *MSGN1-Venus* percent induction as determined by FACS on days 3-4 of differentiation under CL and CLFBR conditions. Mean ±SD. **e,** qRT-PCR comparing relative expression levels of *HES7, LFNG, TBX6, MSGN1, RIPPLY2, MESP2* and *FOXC2* under CL or CLFBR differentiation conditions. Relative expression shown as fold change relative to iPS (day 0). Mean ± SD.

**Extended Data Figure 6.**
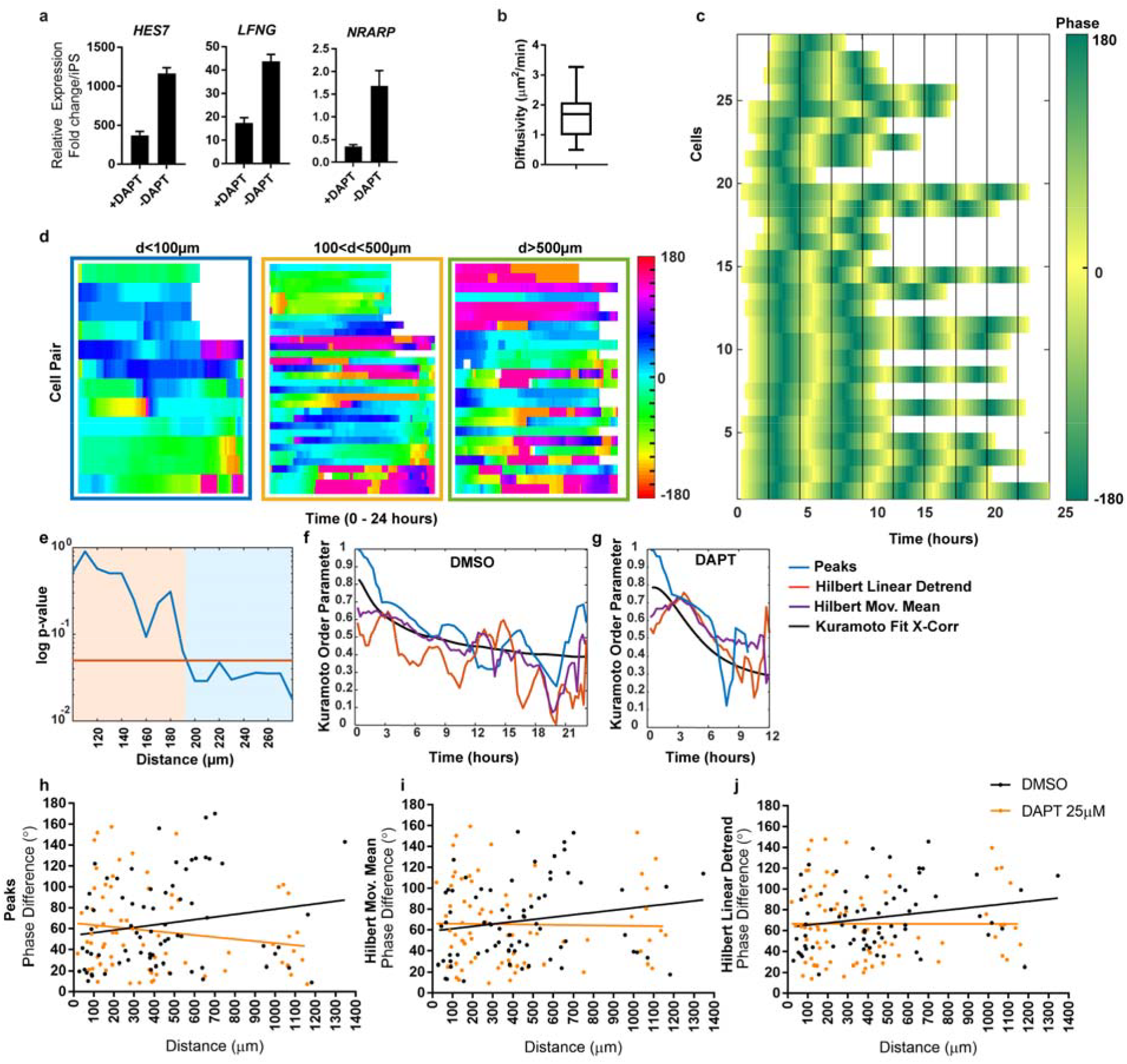
**a,** qRT-PCR for Notch targets *HES7, LFNG* and *NRARP* on day 3 of differentiation (CLFBR medium) in the presence or absence of the γ-secretase inhibitor DAPT (25μM). Mean ±SD **b,** Mean diffusivity (μm^2^/min) for individual *HES7-Achilles* cells over a period of 24 hours. **c,** Evolution of the phase (color-coded) over time for individual cells tracked under control conditions. **d,** Phase shift over time (color-coded) for distinct cell pairs under control conditions, separated into three groups based on the initial distance (μm) between cells. **e,** log p-value as a function of distance, indicating that the difference in average phase shift between nearby and remote cells becomes significant between 190 and 200 μm (interaction distance). **f-g**, Evolution of the Kuramoto order parameter over time in control (f) or 25μM DAPT (g) conditions, calculated using three different definitions of phase. Black line shows the fit based on cross-correlation calculations. **h-j**, Phase difference between pairs of cells as a function of their initial distance in control or 25μM DAPT conditions, calculated with three different definitions of phase: peaks (h), Hilbert moving mean (i), Hilbert linear detrending (j). See methods for details on phase calculations.

**Extended Data Figure 7.**
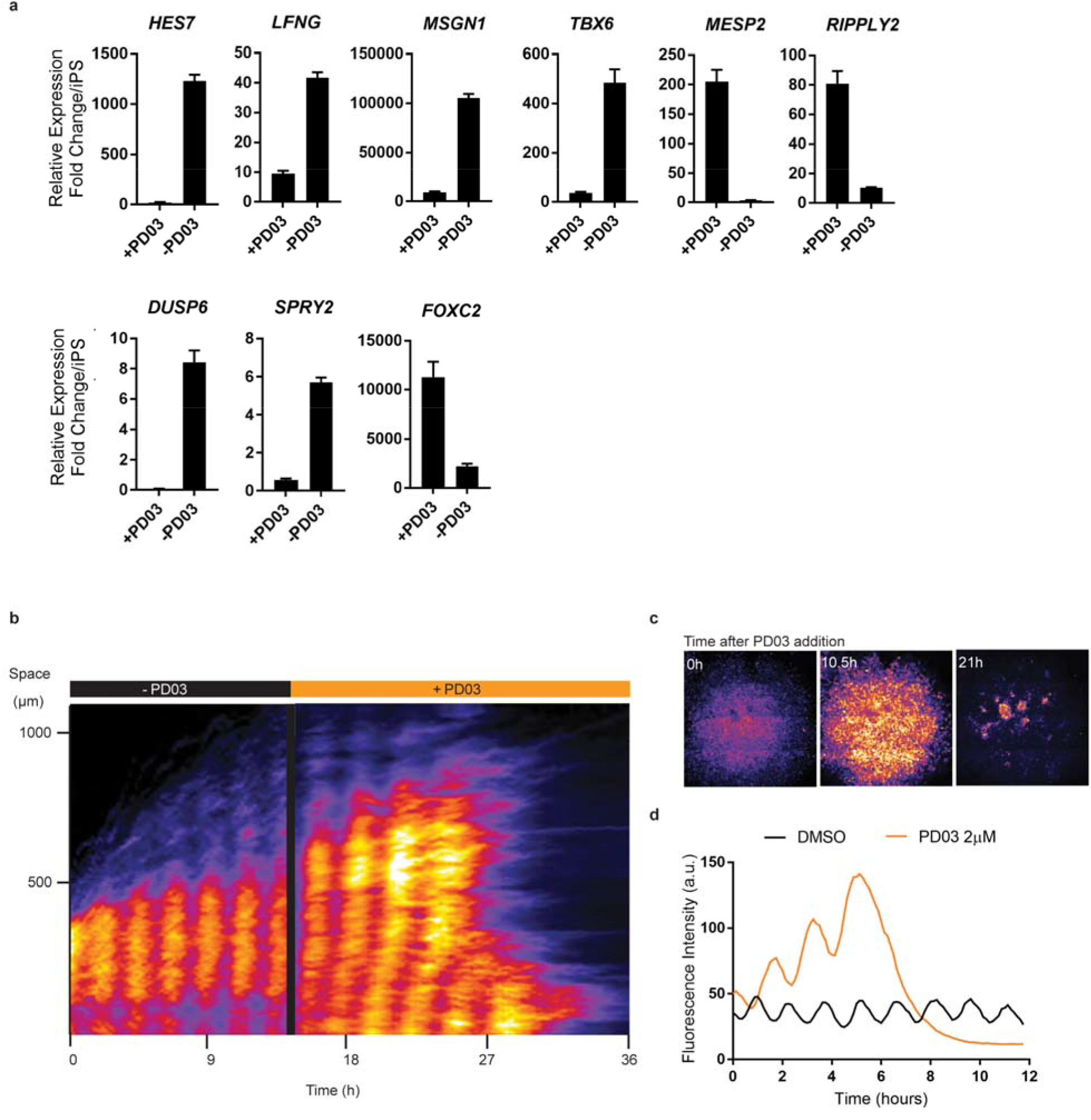
**a,** qRT-PCR comparing relative gene expression levels on day 3 of differentiation (CLFBR medium) in the presence or absence of PD03 (250nM). Relative gene expression shown as fold change relative to iPS (day 0). Mean ±SD **b,** Kymograph of *LuVeLu* oscillations in a mouse tailbud explant before and after the addition of PD03 (2μM). **c,** Snapshots of *LuVeLu* fluorescence in a mouse tailbud explant demonstrating the response to PD03 (2μM) addition. **d,** Quantification of *LuVeLu* fluorescence in mouse tailbud explants treated with either DMSO or PD03 (2μM) over the course of 12 hours.

